# The microbiology and geochemistry of the shallow-water hydrothermal vents of the Gulf of Naples, Italy

**DOI:** 10.1101/2025.10.02.679971

**Authors:** Bernardo Barosa, Carmela Celentano, Flavia Migliaccio, Sara Claudia Diana, Ana Clara Pelliciari Silva, Matteo Selci, Luca Tonietti, Deborah Bastoni, Martina Cascone, Alessia Bastianoni, Monica Correggia, Luciano di Iorio, Roy Price, Stefano Caliro, Marco Milazzo, Alessandro Aiuppa, Costantino Vetriani, Angelina Cordone, Donato Giovannelli

**Author notes:** **Correspondence:** Donato Giovannelli.

## Abstract

Shallow-water hydrothermal vents are dynamic ecosystems that occur below 200 m in tectonically active regions of the planet. While their geochemical composition has been investigated in several locations, knowledge about the microbial diversity they harbour remains scarce. Moreover, the relationships between hydrothermal fluid chemistry, geological settings and microbial community structure in shallow vents have not been explored in detail. Here, we investigate the interplay between fluid geochemistry and microbial diversity in two underwater volcanic regions in the Gulf of Naples, Italy, one under the influence of the Somma-Vesuvio volcano and the other located within the underwater portion of the Campi Flegrei caldera. By combining 16S rRNA amplicon sequencing with geochemical measurements, and by contextualizing it with previous geochemical measurements done in the region, we found that hydrothermal fluid chemistry, influenced by the geological setting where the vents are hosted, plays a key role in shaping microbial ecological niches, and imposes strong selective pressures on the resident microbial communities. We additionally describe two new shallow vent sites, contributing to the characterization of the hydrothermalism in the area and unveiling the biodiversity associated with shallow-water hydrothermalism in the region.

## 1 Introduction

Hydrothermal vents are dynamic marine ecosystems that occur in tectonically active regions of the planet and in proximity to active volcanic regions, where geothermal heat drives fluid circulation in the crust (German and Seyfried, 2014). The resulting hydrothermal fluids enriched in reduced compounds, metals and volatiles mix with oxidised seawater, promoting a redox thermodynamic disequilibrium that fuels highly diverse microbial communities that employ different metabolic strategies (Amend et al., 2003, 2011; Rogers and Amend, 2005, 2006; Dahle et al., 2015, 2018; Price et al., 2015; Rucker et al., 2022). In contrast to their deep-sea counterparts, shallow-water hydrothermal vents (SWHV) occur closer to the surface (below 200 m), and are strongly influenced by solar radiation (Price and Giovannelli, 2017). In these ecosystems chemotrophy and phototrophy co-occur, making SWHVs high energy environments capable of sustaining complex trophic networks (Tarasov et al., 2005; Gomez-Saez et al., 2017; Price and Giovannelli, 2017; Barosa et al., 2023). Due to the shallow depths and consequent decrease in pressure, it is possible to find abundant free-gas phases formed by the exsolution of dissolved volatile gases (Barge and Price, 2022), as well as intense tidal fluctuations and dynamic forcing (Yücel et al., 2013; Price and Giovannelli, 2017; Barge and Price, 2022; Barosa et al., 2023). Additionally, their general proximity to land masses entails a higher transport of terrigenous carbon and phytodetritus into these systems. (Price and Giovannelli, 2017).

Previous studies conducted in shallow-water hydrothermal vents have demonstrated a highly diverse microbial consortia inhabiting these ecosystems (Cardigos et al., 2005; Rusch et al., 2005; Manini et al., 2008; Maugeri et al., 2010, 2013; Giovannelli et al., 2013; Gugliandolo et al., 2015; Sciutteri et al., 2022; Silva et al., 2025b). SWHVs were recently proposed to be a possible cradle for the origin of life or the early evolution of microbial metabolism (Barge and Price, 2022; Rucker et al., 2022). Thermodynamic mapping of exergonic reactions has demonstrated a high energy landscape available for microbial metabolism (Amend et al., 2003; Rogers and Amend, 2005, 2006; Rogers et al., 2007; Price et al., 2015; Lu et al., 2020). Depending on the typology of the hydrothermal system, niche stratification can be observed, with a shift from sites dominated by chemolithoautotrophic and sulfur utilising microbial communities to an increase of phototrophic and heterotrophic communities (Giovannelli et al., 2013; Gugliandolo et al., 2015; Fagorzi et al., 2019; Barosa et al., 2023; Silva et al., 2025b).

Differences in fluid geochemistry and the geological setting where hydrothermal systems are hosted have been shown to impose strong controls on the resident microbial communities (Fullerton et al., 2021; Colman et al., 2023; Sims et al., 2023; Upin et al., 2023). As water percolates into the crust, fluids are transformed by subsurface processes, such as temperature dependent water-rock-gas reactions, the release of metals and other elements from the host lithologies, as well as phase separation events (Tivey, 2007; German and Seyfried, 2014; Price et al., 2015). As such, the composition of each hydrothermal fluid can be, to some degree, unique to every hydrothermal system, reflecting changes in rock type, shape and size of the heat sources, composition of the primary volatiles, geochemical reactions happening during fluid ascent, timescales of fluid-rock interaction (German and Seyfried, 2014; Ely et al., 2023). The typology of the system also exerts control on how the fluids are being discharged at the surface. For instance, hydrothermal systems hosted close to a heat source (e.g. magmatic chamber or intrusions) or with direct channelling to the surface, usually present higher temperatures and more focused flows, compared to more diffuse fluids, where there is a higher degree of mixing between the hydrothermal fluids with seawater (Von Damm and Lilley, 2004; Scheirer et al., 2006). This mixing also constrains the possible niches occupied by microbial communities, given the variance in electron donors/acceptors available, as well as different carbon sources. However, the extent to which these different geochemical landscapes constrain shallow-water hydrothermal vents communities has not been explored in detail.

A previous study conducted in SWHV of the Aeolian archipelago (Italy) reported that microbial communities changed according to the different geochemical regimes found within the different islands, following large-scale trends in volcanic activity (Barosa et al., 2023). Here, we combine 16S rRNA amplicon sequencing with geochemical and physico-chemical measurements of the shallow-water hydrothermal vents present within two volcanic regions of the Gulf of Naples, in the Campania region, Italy, to investigate how the geochemistry and typology of the hydrothermal systems in the area structure the microbial communities. The Gulf of Naples is characterized by the presence of active volcanisms, with three active volcanoes in the area that are either surrounded or extend at sea. Given the recent signs of unrest with increasing seismic activity of the Campi Flegrei volcano (Astort et al., 2024; Giudicepietro et al., 2025), this study may provide baseline microbiological data for future research aimed at understanding shifts in microbial communities in response to changes in volcanic activity.

## 2 Experimental Procedures

### 2.1 Campi Flegrei and Somma-Vesuvius volcanic complexes

Naples’s area is characterized by the presence of three active volcanoes: the Ischia volcanic complex, the Campi Flegrei caldera and the Somma-Vesuvius volcanic complex. The gulf is characterized by intense hydrothermal activity, with numerous submarine hydrothermal systems reported in the region (Figure 1) (De Astis et al., 2004; Aiello et al., 2010; Kerrison et al., 2011; Calosi et al., 2013; Torrente and Milia, 2013; Passaro et al., 2014b; Somma et al., 2016; Donnarumma et al., 2019; Baldrighi et al., 2020; Bellec et al., 2020; Sacchi et al., 2020).

**Figure 1.**
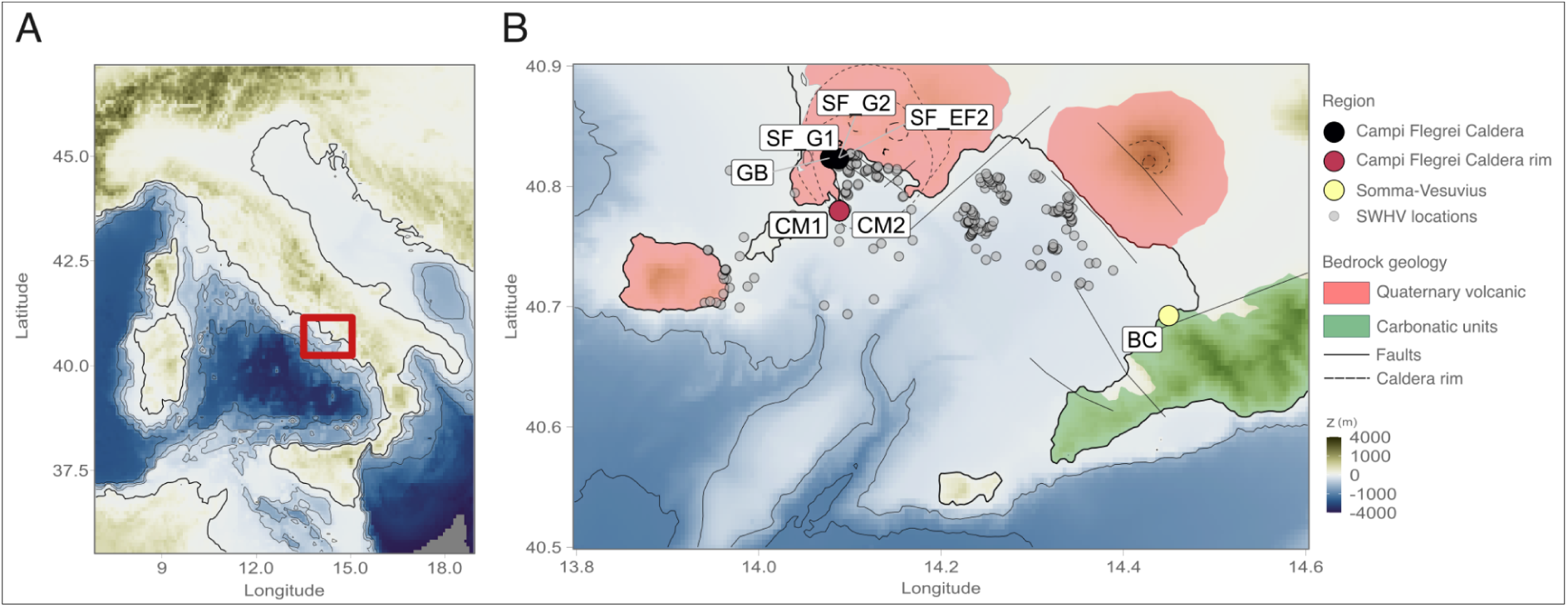
Map depicting the shallow-water hydrothermal vents sampled. **A -** Regional map of Italy (red square indicating the sampled locations). **B -** Map of the Gulf of Naples. Coloured points represent the different locations sampled in the present study, along the bathymetry depiction of the region. Grey points are known shallow-water hydrothermal vents from the literature (De Astis et al., 2004; Aiello et al., 2010; Kerrison et al., 2011; Calosi et al., 2013; Torrente and Milia, 2013; Passaro et al., 2014b; Somma et al., 2016; Donnarumma et al., 2019; Baldrighi et al., 2020; Bellec et al., 2020; Sacchi et al., 2020).

The Campi Flegrei caldera is a significant and historically active volcanic field situated along the Neapolitan Tyrrhenian coastline of Italy. The distinctive geomorphology of the Pozzuoli Bay area has been shaped by two major explosive eruptions: the Campanian Ignimbrite approximately 39,000 years ago (Rosi et al., 1996) and the Neapolitan Yellow Tuff around 15,000 years ago (Orsi et al., 1992). The southern part of the caldera is now submerged (the Pozzuoli Bay area), and is hence characterised by intense underwater hydrothermal activity. The area has been characterized by uplifting and intense seismic events (Astort et al., 2024; Giudicepietro et al., 2025), and the inland and submarine geothermal features are constantly monitored to understand the state of unrest of the volcano. The last eruption is dated 1538, with the formation of Monte Nuovo.

On the western side of the city of Naples, the Somma-Vesuvius volcanic complex originated from a sequence of eruptions through time, starting from the construction of the Somma edifice (>20,000 years ago) that was later (partially) dismantled in a sequence of Plinian eruptions between 20,000 to 1,871 years ago. These eruptions gave rise to the formation of a summit caldera, measuring 4.9 by 3.4 kilometres in an east-west elongated configuration (Bertagnini et al., 1998), topped by the young (post-caldera) Vesuvius cone. Mild hydrothermal activity has been ongoing at Vesuvius since its last eruption in 1944. Key indicators of this activity include: (a) weak fumarolic emissions, accompanied by diffuse soil CO_2_ degassing in the crater area (Chiodini et al., 2001; Frondini et al., 2004); (b) CO_2_-rich groundwaters along the southern flank of Vesuvius and in the adjacent plain (Caliro et al., 2011); and (c) episodic seismic activity with epicentres concentrated within the crater (Caliro et al., 2011). The toe-shaped morphology on the seafloor located in the southwestern side of the Somma-Vesuvius is explained by two flank collapses in the Late Pleistocene (Passaro et al., 2014a).

### 2.2 Sampling and site description

Samples were collected from June to October 2020 during the FEAMP20 expedition on the shallow-water hydrothermal vents present on the Gulf of Naples (Figure 1) following a large-scale sampling approach (Giovannelli et al., 2022). More specifically, samples were collected at the region of Campi Flegrei caldera: Secca delle Fumose (SF) and Gabbiano (GB), at the region of Campi Flegrei caldera rim: Capo Miseno (CM), and the region of Somma-Vesuvius: Bagno Conte (BC). The most interesting sites, Secca delle Fumose (geyser) and Capo Miseno, were sampled twice, and indicated as SF_G2 and CM2, respectively (Table 1). All samples were retrieved by scuba divers, and in order to get a broader characterization of the vent environment, three types of samples were collected at each site: sediments, fluids, and biofilms. Fluids were sampled through the insertion of a silicon tube in the selected vent orifice (7-15 cm), and filtered through Sterivex 0.22 µm filter membranes. The sediments were collected in the vent orifice (top 1-2 cm), and biofilms were collected at or in the vicinity of the vents. For geochemical analyses, fluids were filtered (0.22 µm) and treated with nitric acid (final concentration of 2 %). In the laboratory, Sterivex filters and sediments were stored at -20 °C until DNA extraction, while the samples for geochemical analyses were stored at 4 °C. Physico-chemical parameters for each sample site were measured *in-situ* using a multiparametric probe (HANNA, HI98196), a refractometer (Fisherbrand ™), and a field thermometer.

**Table 1.**
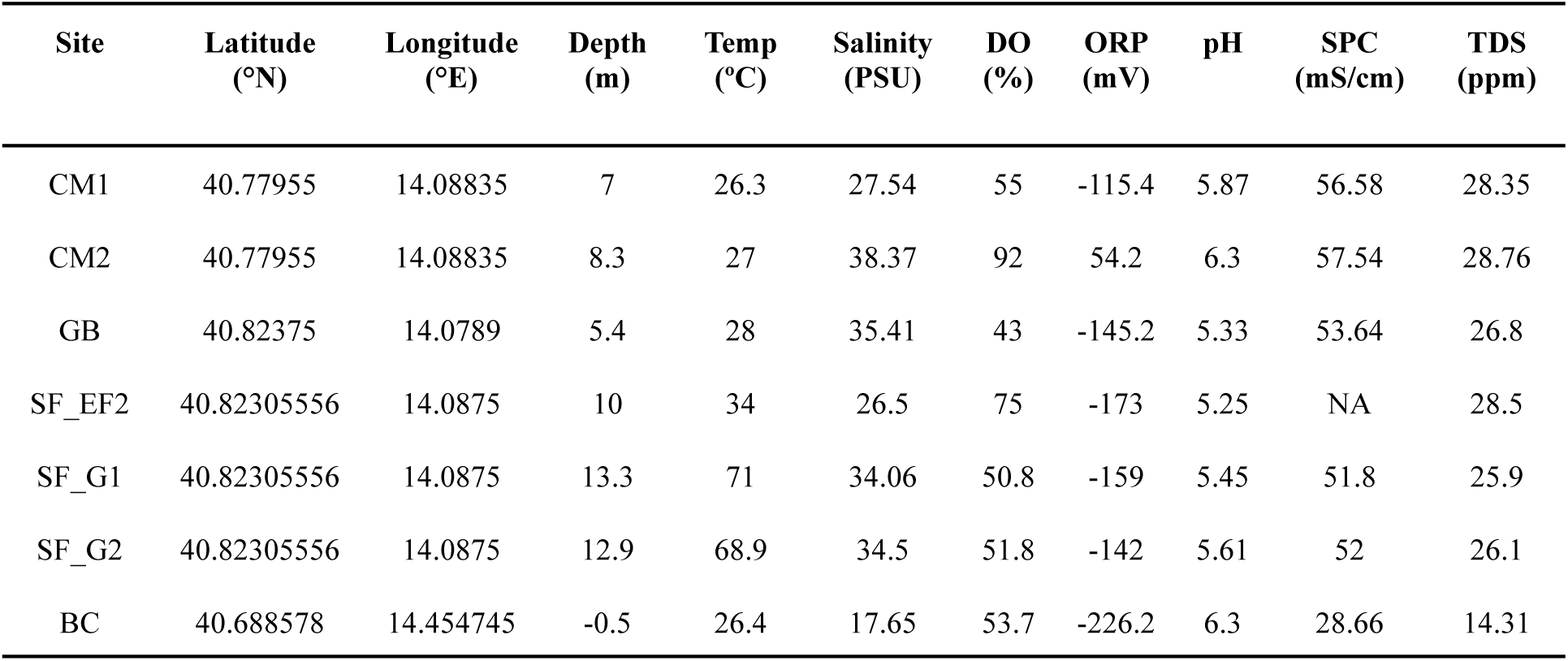
Location (GPS coordinates), Depth, and the Physico-chemical parameters measured on the venting fluids at the different sampled locations. ORP - Oxido-reduction potential; DO - Dissolved oxygen; mS/cm - Conductivity; ppt tds - Total dissolved solids.

| Site | Latitude<br>(°N) | Longitude<br>(°E) | Depth<br>(m) | Temp<br>(°C) | Salinity<br>(PSU) | DO<br>(%) | ORP<br>(mV) | pH | SPC<br>(mS/cm) | TDS<br>(ppm) |
| --- | --- | --- | --- | --- | --- | --- | --- | --- | --- | --- |
| CM1 | 40.77955 | 14.08835 | 7 | 26.3 | 27.54 | 55 | -115.4 | 5.87 | 56.58 | 28.35 |
| CM2 | 40.77955 | 14.08835 | 8.3 | 27 | 38.37 | 92 | 54.2 | 6.3 | 57.54 | 28.76 |
| GB | 40.82375 | 14.0789 | 5.4 | 28 | 35.41 | 43 | -145.2 | 5.33 | 53.64 | 26.8 |
| SF_EF2 | 40.82305556 | 14.0875 | 10 | 34 | 26.5 | 75 | -173 | 5.25 | NA | 28.5 |
| SF_G1 | 40.82305556 | 14.0875 | 13.3 | 71 | 34.06 | 50.8 | -159 | 5.45 | 51.8 | 25.9 |
| SF_G2 | 40.82305556 | 14.0875 | 12.9 | 68.9 | 34.5 | 51.8 | -142 | 5.61 | 52 | 26.1 |
| BC | 40.688578 | 14.454745 | -0.5 | 26.4 | 17.65 | 53.7 | -226.2 | 6.3 | 28.66 | 14.31 |

### 2.3 Geochemical Analysis

The concentrations of major cations (sodium-Na^+^, potassium-K^+^, magnesium-Mg^2+^, ammonia-NH_4_ ^+^, and calcium-Ca^+^) and anions (chlorine-Cl^-^, sulfate-SO_4_ ^2-^ and bromide-Br^-^) of the sampled fluids were measured (in duplicates) using ion chromatography (ECO, IC Metrohm). Calibration curves for the ion species were measured in the range of 0.1 and 200 ppm with an R ≥ 0.999. All samples were filtered (0.22 μm) and diluted to a conductivity of 600 µS/cm (Correggia et al., 2023a). All dilutions were made using 18.2 MΩ/cm type I water, also used as a blank for blank subtractions. Anions were run using a 3.2 mM Na_2_CO_3_ + 1 mM NaHCO_3_ mobile phase on a Metrosep A Supp 5 column equipped with a 0.15 M ortho-Phosphoric acid suppressor. The flow of the anionic eluent was 0.7 ml min^-1^ for 30 min. Cations were run using a 2.5 mM HNO_3_ + 0.5 mM (COOH)_2_ × 2H_2_O mobile phase on a Metrosep C4 column. The flow of the cationic eluent was 0.9 ml min^-1^ with a total separation of 35 min. Data acquisition and analysis were carried out through MagIC Net 3.3 software. Calibration curves were defined using certified CPA chem external multi-ion standards for each of the anions and cations analysed in this study (1000 mg/L). Additionally, the concentration of biologically relevant trace elements (i.e. Fe, Zn, As, Rb, Sr, Mo and Cs), were measured (in triplicates) using an Agilent inductively coupled plasma mass spectrometer (Agilent ICP-MS 7900) following the standard operating practice developed at the Giovannelli Lab (Correggia et al., 2023b). Calibration curves covered a concentration range between 0.01 to 100 ppb, and were run using Agilent certified multistandard 2A (10 μg/mL of Ag, AI, As, Ba, Be, Ca, Cd, Co, Cr, Cs, Cu, Fe, Ga, K, Li, Mg, Mn, Na, Ni, Pb, Rb, Se, Sr, TI, U, V, Zn). Additionally, we contextualised the geochemistry of these hydrothermal systems with published data from the region found in the literature. The visualisation and statistical analysis on geochemical data were carried out in the R statistical software version 4.1.2 (Core, 2021) and ggplot2 packages (Wickham, 2011).

### 2.4 DNA Extraction and Amplicon Sequencing

DNA extraction was performed using the DNeasy PowerSoil Kit (QIAGEN), following the manufacturer’s instructions, with an extra elution step. When low DNA yields were obtained, a modified phenol-chloroform DNA extraction method was used, adapted for shallow-water hydrothermal vent conditions (Giovannelli et al., 2013, 2016b). Briefly, 850 µl of extraction buffer (50 mM Tris-HCl pH 8, 20 mM EDTA pH 8, 100 mM NaCl) was added to 0.5 g of sediment and sterivex filters. Followed with 100 µl of Lysozyme (100 mg/ml) and 5 µl of Proteinase K, with a step of incubation at 37 °C for 30 min after each supplementation step, and followed by 1 min of vortexing. Subsequently, 50 µl of 20 % SDS was added to the samples and incubated for 30 min at 65 °C, with regular mixing, followed by a vortexing step for 1 min. Samples were then centrifuged for 3 min at 14,000 × g. The supernatant was collected and transferred to a phenol:chloroform:isoamyl alcohol (25:24:1) solution, followed by centrifugation for 3 min at 14,000 × g. The aqueous phase was collected and transferred to a solution of chloroform:isoamyl alcohol (24:1) and centrifuged for 3 min at 14,000 × g. The supernatant was collected and supplemented with Na-acetate (0.1 vol) and isopropanol (0.7 vol), and incubated at -20 °C for 2 h. The precipitated DNA was further washed with 70% cold ethanol and re-suspended in 50 μl of Tris-HCL. DNA was visualised with agarose gel electrophoresis and quantified spectrophotometrically and spectrofluorimetrically (NanoDrop and Qubit, respectively). The successfully extracted DNA was sequenced at the Integrated Microbiome Resource (IMR, https://imr.bio) using primers targeting the V4-V5 region (515FB = GTGYCAGCMGCCGCGGTAA and 926R = CCGYCAATTYMTTTRAGTTT515FB - 926R) of the 16S rRNA gene, using an Illumina V3 sequencing MiSeq.

### 2.5 Bioinformatic and statistical analysis

The raw sequencing data were analysed using the DADA2 package (Callahan et al., 2017). Quality profile analysis was carried out after trimming the primers and adapters, where only the sequences with a quality call for each base between 20 and 40 were kept for downstream analysis. Amplicon sequencing variants (ASVs) were estimated through error profiling, and taxonomy was assigned to each ASV with the SILVA database, release 138 (Quast et al., 2013). Amplicon variant abundance tables and taxonomic assignments were used to create a phyloseq object to further calculate diversity indices and investigate the microbial diversity, as previously reported (Cordone et al., 2022, 2023; Barosa et al., 2023). Sequences belonging to Mitochondria, Eukaryotes, Chloroplasts, groups related to human pathogens and common DNA extraction contaminants (Sheik et al., 2018), as well as the sequenced blanks, were removed from the dataset. The remaining reads represented 95 % of the original reads, with 178671 reads assigned to 4612 ASVs. Diversity analyses were carried out using the Phyloseq package (McMurdie and Holmes, 2013) with the relative abundance values set to 100 %. Beta diversity was investigated using the Jaccard diversity index (Unweighted and Weighted) as implemented in the vegan package (Oksanen et al., 2018). The obtained ordination was used to investigate the correlation between the environmental and geochemical variables using environmental fitting (*envfit* and *ordisurf* functions). ASV taxonomic classification was further carried out using the EzBioCloud database (Yoon et al., 2017), as well as blasted against the nucleotide database of the National Center for Biotechnology Information (NCBI). All statistical calculations, data processing, and visualisation were carried out in the R statistical software version 4.1.2 (Core, 2021) and ggplot2 packages (Wickham, 2011). All the sequences analysed in this study are available through the European Nucleotide Archive (ENA) under project accession PRJEB67762, under the ENA Umbrella project CoEvolve PRJEB55081. A complete R script containing all the steps of our analysis is available at https://github.com/giovannellilab/Barosa_et_al_Feamp20_16S with DOI: 10.5281/zenodo.14966493, together with all the environmental and geochemical data.

## 3 Results

### 3.1 Site description and physico-chemical parameters

The samples were collected in three different locations belonging to two different volcanic regions (Figure 1), the Campi Flegrei caldera (SF_G1, SF_G2, SF_EF, and GB) and caldera rim (CM1 and CM2), and the Somma-Vesuvius area (BC). Venting in the SF_G1 site, known by local divers as the geyser, occurs in a rocky bottom and it is characterized by intense focused emission surrounded by yellow, orange and white bacterial mats and mineral precipitates. No free gas is visible at this site. Venting in the SF_EF site, located in close proximity to SF_G1, occurs in a field of large rocks . Gas bubbles are visible and run against the rocks determining the distribution of abundant white biofilms. Venting at site GB is diffuse and occurs through sediments covered with white particles. Minimal free gas is present in the area. The site CM, located in the Campi Flegrei caldera rims is characterized by diffuse gas emission in a rocky area and the presence of a larger more focused opening named “the flute”, covered with white filamentous biofilms. The site BC, in the region of Somma-Vesuvius, is located at the shoreline, and characterised by white particles in suspension and an intense smell of “rotten eggs” characteristic of H_2_S.

The physico-chemical parameters of the shallow-water hydrothermal vents of the Gulf of Naples sampled in this study are shown in Table 1. Vent temperatures varied from 26.3 °C in the SWHV of CM1 to 71 °C in SF_G1. The salinity ranged from 17.65 PSU in BC to 38.37 PSU in CM2. Dissolved oxygen (DO) values ranged from 43 % in GB to 92 % in CM2. The oxidation-reduction potential (ORP) was negative for all the sampling sites with the exception of CM2, in which the value is 54.2 mV. The pH was slightly acidic for every sampled shallow-water vent (average pH of 5.73). The conductivity for each site is around 50 mS/cm, with the exception of BC, in which the value is 28.7 mS/cm. The total dissolved solid (TDS) ranged between 14.31 ppt TDS in BC to 28.76 ppt TDS in CM2.

### 3.2 Geochemical Results

The concentrations of the major and minor elements (and respective measurement errors reported in % RSD) of the shallow-water hydrothermal vents sampled in this study are presented in Table 2 and Table 3, respectively. Chloride (Cl^-^) ranged from 10,030.06 ppm in the BC site to 21,094.04 ppm at CM1. Bromide (Br^-^) values ranged from 34.27 ppm in BC to 72.48 ppm in CM1, whereas sulfate (SO_4_ ^2-^) ranged from 1,377.55 ppm in BC to 2,920.13 ppm in SF_EF. Sodium (Na^+^) values ranged from 5,844.84 ppm in BC to 12,783.86 ppm in SF_EF, while ammonium (NH ^+^) ranged from 33.28 ppm in GB to 40.30 ppm in SF_G2. Potassium (K^+^), magnesium (Mg^2+^), and calcium (Ca^2+^) concentrations ranged from 198.47 ppm in BC to 598.43 ppm in GB, 606.89 ppm in BC to 1,395.17 ppm in CM1, and 434.79 ppm in SF_EF to 497.28 ppm in SF_G2, respectively.

**Table 2.**
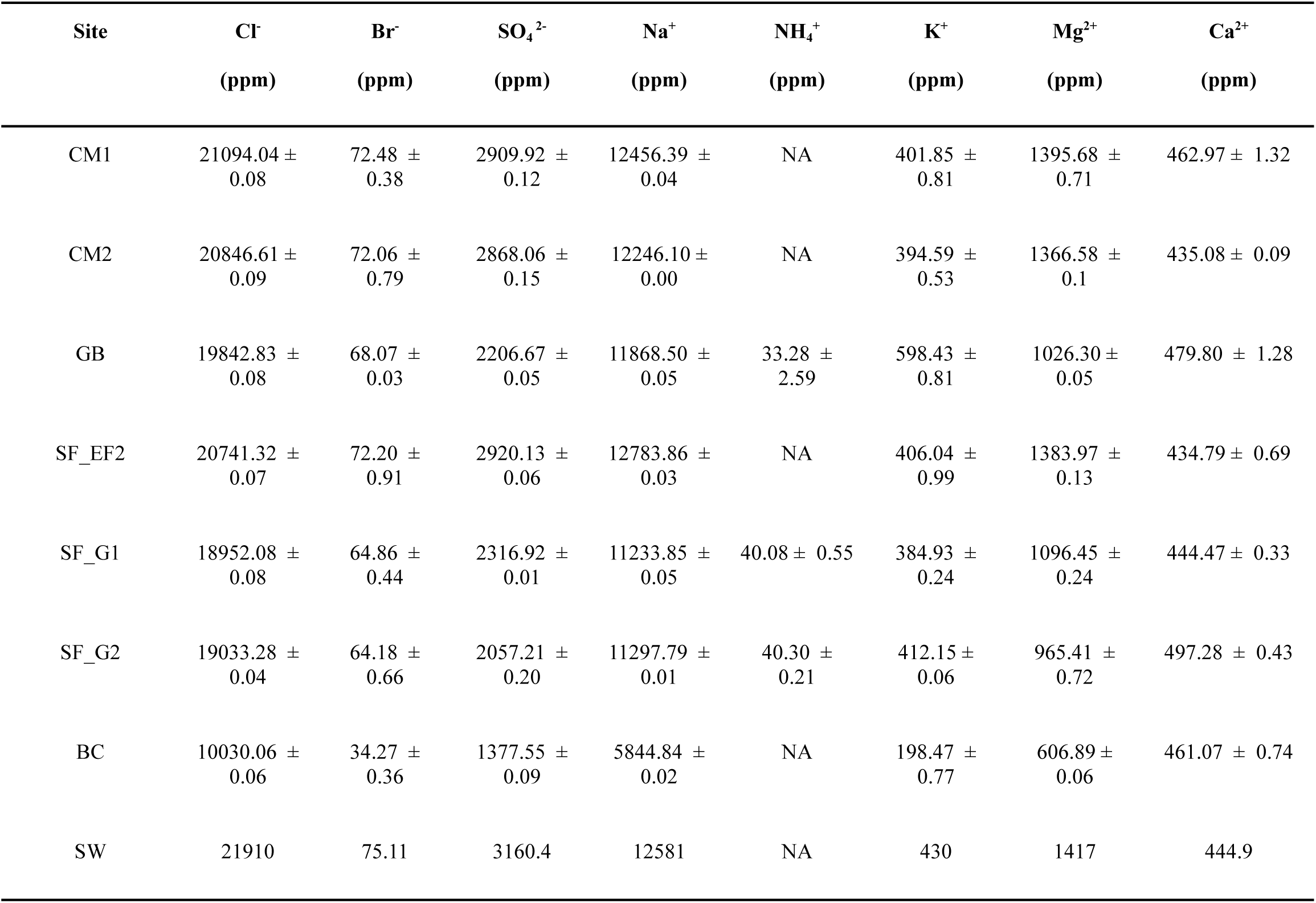
Concentrations (ppm) of the major anions and cations of the different sampled shallow-water hydrothermal vents (error ± in % RSD). Seawater (SW) values were obtained from Price et al., 2015.

**Table 3.**
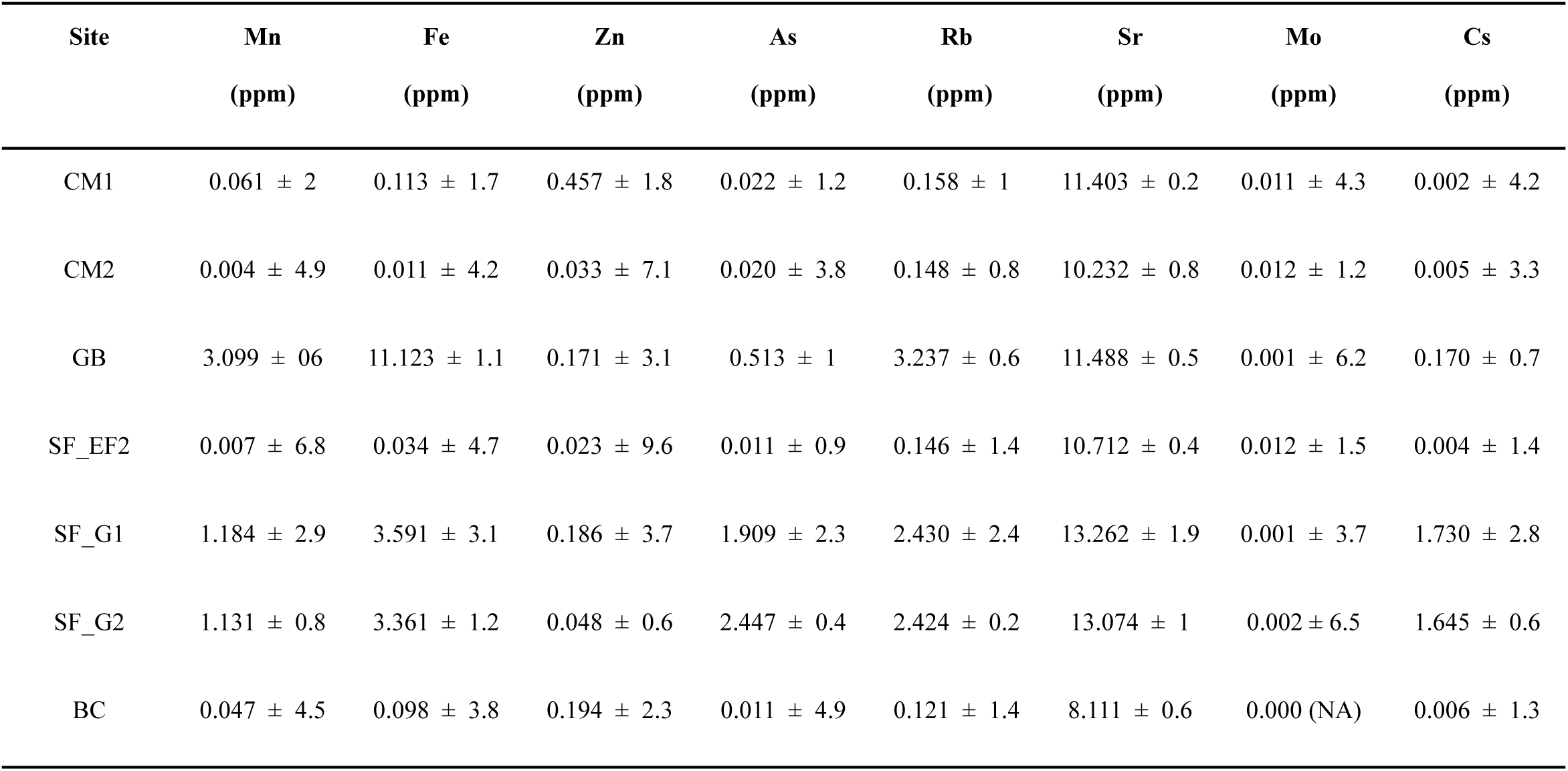
Concentrations (ppm) of the trace elements of the different sampled shallow-water hydrothermal vents (error ± in % RSD).

| Site | Mn<br>(ppm) | Fe<br>(ppm) | Zn<br>(ppm) | As<br>(ppm) | Rb<br>(ppm) | Sr<br>(ppm) | Mo<br>(ppm) | Cs<br>(ppm) |
| --- | --- | --- | --- | --- | --- | --- | --- | --- |
| CM1 | 0.061 ± 2 | 0.113 ± 1.7 | 0.457 ± 1.8 | 0.022 ± 1.2 | 0.158 ± 1 | 11.403 ± 0.2 | 0.011 ± 4.3 | 0.002 ± 4.2 |
| CM2 | 0.004 ± 4.9 | 0.011 ± 4.2 | 0.033 ± 7.1 | 0.020 ± 3.8 | 0.148 ± 0.8 | 10.232 ± 0.8 | 0.012 ± 1.2 | 0.005 ± 3.3 |
| GB | 3.099 ± 06 | 11.123 ± 1.1 | 0.171 ± 3.1 | 0.513 ± 1 | 3.237 ± 0.6 | 11.488 ± 0.5 | 0.001 ± 6.2 | 0.170 ± 0.7 |
| SF_EF2 | 0.007 ± 6.8 | 0.034 ± 4.7 | 0.023 ± 9.6 | 0.011 ± 0.9 | 0.146 ± 1.4 | 10.712 ± 0.4 | 0.012 ± 1.5 | 0.004 ± 1.4 |
| SF_G1 | 1.184 ± 2.9 | 3.591 ± 3.1 | 0.186 ± 3.7 | 1.909 ± 2.3 | 2.430 ± 2.4 | 13.262 ± 1.9 | 0.001 ± 3.7 | 1.730 ± 2.8 |
| SF_G2 | 1.131 ± 0.8 | 3.361 ± 1.2 | 0.048 ± 0.6 | 2.447 ± 0.4 | 2.424 ± 0.2 | 13.074 ± 1 | 0.002 ± 6.5 | 1.645 ± 0.6 |
| BC | 0.047 ± 4.5 | 0.098 ± 3.8 | 0.194 ± 2.3 | 0.011 ± 4.9 | 0.121 ± 1.4 | 8.111 ± 0.6 | 0.000 (NA) | 0.006 ± 1.3 |

The trace elements measured in this study are presented in Table 3. Manganese (Mn) concentrations ranged from 0.0037 ppm in CM2 to 3.1 ppm in GB, while iron (Fe) concentrations ranged from 0.011 ppm in CM2 to 11.12 ppm in GB. Zinc (Zn) and Arsenic (As) concentrations ranged from 0.023 ppm in SF_EF2 to 0.46 ppm in CM1, and 0.011 ppm in SF_EF2 to 2.45 ppm in SF_G2 in SF_G2, respectively. Rubidium (Rb) concentrations ranged from 0.12 ppm in BC to 3.23 ppm in GB. Strontium (Sr) and Caesium (Cs) values ranged from 8.11 ppm in BC to 13.3 in SF_G1, and 0.002 ppm in CM1 to 1.73 ppm in SF_G1, respectively. Molybdenum (Mo) concentrations ranged from 0 ppm in BC to 0.012 ppm in SF_EF2. Seawater element concentrations were obtained from Price et. al., 2015.

Principal Component Analysis (PCA, Figure 2A) constructed using the physico-chemical characteristic of each site and major ion composition shows that sites clustering according to the geographic locations where venting occurs, with the exception of SF_EF. Site BC, located under the influence of the Somma-Vesuvius volcanic complex, presented the most diverse geochemistry compared to all the other sites investigated and clustered on its own (Figure 2A). The concentrations of chloride plotted against sodium suggest that all the fluids are of seawater origin and mixed with lower salinity deeply-derived fluids with the exception of BC, that might be the results of meteoric recharge mixing with seawater and shows much lower salinities (Figure 2B). The mixing plots of the other major ions (Figure 2D–E) support this trend, revealing three distinct clusters (consistent with the PCA results). Sites SF_EF, CM1, and CM2 exhibit concentrations similar to seawater, indicating a higher degree of fluid mixing with background seawater, compared to sites SF_G1, SF_G2, and GB. The higher Fe concentrations observed at site GB (Figure 2F) suggest increased rock leaching, possibly influenced by the local host lithology.

**Figure 2.**
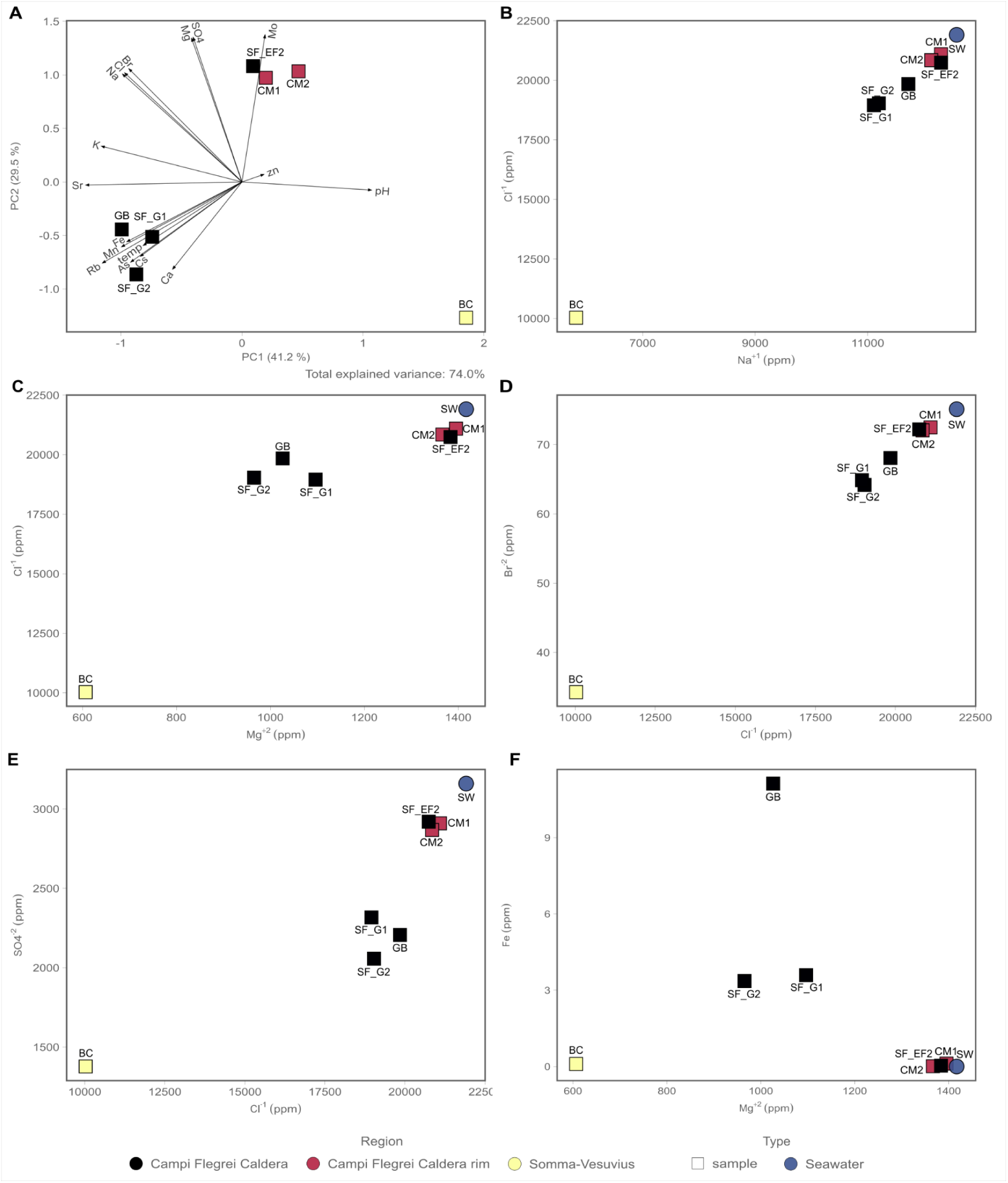
Geochemical data measured at the different shallow-water hydrothermal vents of the Campi Flegrei region and Somma-Vesuvius. Seawater concentrations were taken from Price et al., 2015**. A)** Principal component analysis (PCA) with geochemical data measured (major ions and trace elements; see Table 2 and 3). **B)** Biplot Cl^-^ against Na^+^ **C)** Biplot Cl^-^ against Mg^2+^ **D)** Biplot Br^2-^ against Cl^-^ **E)** Biplot SO_4_^2-^ against Cl^-^ and **F)** Biplot Fe against Mg^2+^. Values reported in ppm and errors related to the measurements are reported in Table 2 and 3.

### 3.3 Microbial Diversity

The microbial diversity of the shallow-water vents of the Gulf of Naples was investigated using the V4-V5 regions of the 16S rRNA gene. A total of 4,612 ASVs were identified (after quality check and filtering steps, see methods for details).

The sites SF_G1, SF_G2, SF_EF, and GB are in the Campi Flegrei caldera region (Figure 3). The site SF_G1, represented only by a fluid sample, was dominated by the Aquificota phylum, followed by Chloroflexi, Desulfurobacterota, and an unidentified phylum, at lower abundances. Within Aquificota, *Hydrogenothermus* was the most abundant genus in this site (63.0 % average relative abundance). Within Chloroflexi, the most abundant family was *Anaerolineaceae*, with no classification down to the genus level (8.8 % average relative abundance). Within Desulfurobacterota, the most abundant genera belonged to the *Thermodesulforhabdus* and *Thermosulfurimonas* (4.5 % and 3.0 % average relative abundance, respectively). An unidentified phylum was also present in lower abundances (2.1 % average relative abundance). The site SF_G2, composed of both fluid and sediment samples, was dominated by members belonging to the phyla Proteobacteria, Cyanobacteria and Bacteroidota, at lower abundances. Within Proteobacteria, the most abundant family belonged to *Thiomicrospiraceae*, with no further classification down to the genus level (5.2 % average relative abundance across fluid and sediment samples), followed by a gammaproteobacterial unidentified ASV (3.0 % average relative abundance across fluid and sediment samples). Additionally, it was possible to identify the genera *Thiogranum*, and *Woeseia*, the Sars11 clade, and the genus *Thiohalophilus* (2.4 %, 2.3 %, and 2.2 % average relative abundance across fluid and sediment samples, respectively). Within Cyanobacteria, the most abundant genus belonged to *Synechococcus* CC9902 (3.6 % average relative abundance across fluid and sediment samples). Within Bacteroidota, the most abundant family belonged to *Cryomorphaceae*, with no further taxonomic classification (2.2 % average relative abundance across fluid and sediment samples).

**Figure 3.**
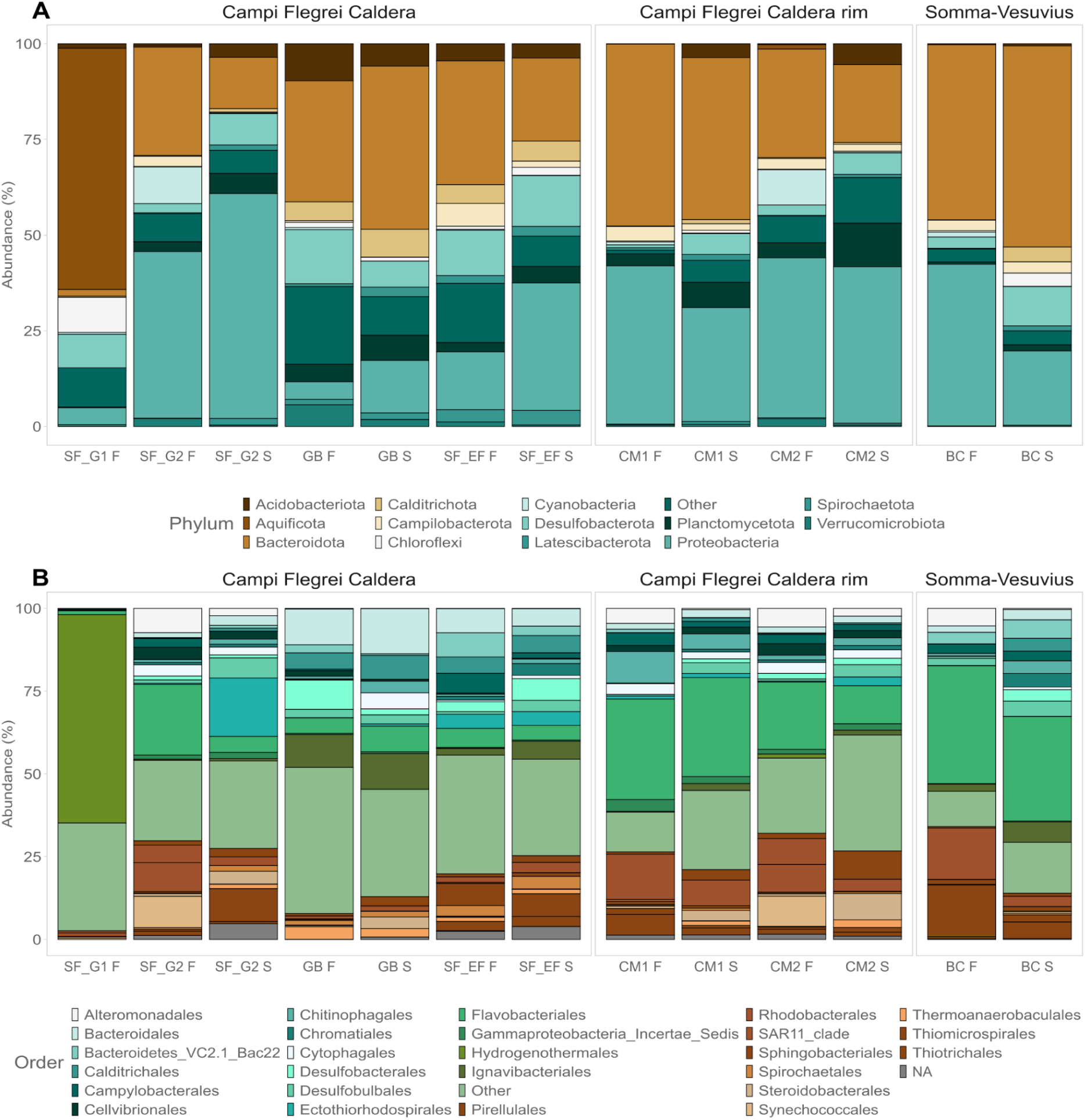
Microbial community composition of the different shallow-water hydrothermal vents sampled, with the two matrices, sediments and fluids. A) Phylum taxonomic rank; B) Order taxonomic rank.

The site SF_EF has as most abundant members, ASVs affiliated to the Bacteroidota phylum, with Calditrichota, Desulfurobacterota, Halobacterota, and Campylobacterota, present in lower abundances. Within Bacteroidota, the most abundant ASV belongs to the clade bacteroidetes VC2.1 Bac22 (5.0 % average relative abundance across the fluid and sediment samples), and to the families *Lentimicrobiaceae*, *Melioribacteraceae*, and *Bacteroidetes* BD2-2 (3.8 %, 3.1 %, and 2.1 % average relative abundance across fluid and sediment samples, respectively). Within Calditrichota, Desulfurobacterota, Halobacterota, and Campylobacterota, ASVs were affiliated with the genera *Caldithrix*, *Desulfovibrio*, *Methanofolis*, and *Sulforovum*, respectively (3.3 %, 2.7 %, 2.7 %, 2.5% average relative abundance across fluid and sediment samples, respectively).

The site GB, located near both SF_G1 and SF_G2, is the one in this region closest to land. This site is composed of both fluid and sediment samples, and dominated by members belonging to the Bacteroidota phylum, with Acidobacterota, Desulfurobacterota, Nanoarchaeota, and an unidentified phylum, present at lower abundances. Within Bacteroidota, the most abundant genus belongs to *Marinifilum* (7.6 % average relative abundance across fluid and sediment samples), followed by the clade PHOS-HE36 and family *Cyclobacteriaceae*, which could not be resolved to the genus level (5.9 % and 2.5 % average relative abundance across fluid and sediment samples, respectively). Additionally, it was possible to identify the clade lheB3-7 (2.2 % average relative abundance across fluid and sediment sample, as well as an ASV belonging to an unidentified phylum (3.9 % average relative abundance across fluid and sediment sample). Within Acidobacteriota, the genus *Thermotomaculum* was present in higher abundances (2.5 % average relative abundance across fluid and sediment samples). Within Desulfurobacteriota and Nanoarchaeota, it was possible to identify the family *Desulfosarcinaceae* and the clade SCGC_AAA011-D5, with no further taxonomic classification (2.1% average relative abundance in both the fluid and sediment samples).

The sites CM1 and CM2 are located within the Campi Flegrei caldera rim region. The site CM1, composed of both fluid and sediment samples, was dominated by the phylum Bacteroidota and Proteobacteria, with Campylobacterota present at lower abundances. Within Bacteroidota, the most abundant family belonged to the *Flavobacteriaceae*, where it was possible to identify the genera *Lutibacter*, *Olleya*, and *Maritimimonas* (8.0 %, 3.7 %, and 2.0 % average relative abundance across fluid and sediment samples, respectively). Additionally, one ASV could not be resolved to the genus level (5.1 % average relative abundance in both the fluid and sediment samples). The ASVs belonging to the families *Saprospiraceae* and *Cryomorphaceae* were not possible to classify down to the genus level (4.2 % and 2.6 % average relative abundance across fluid and sediment samples). Within Proteobacteria, the most abundant family belonged to the *Rhodobacteraceae*, where one ASV could not be classified to the genus level, and the other to the genus *Actibacterium* (3.8 % and 2.1 % average relative abundance across fluid and sediment samples). The family *Thiotrichaceae* could not be resolved to the genus level (3.0 % average relative abundance across fluid and sediment samples). Within Campylobacterota, *Sulfurovum* was the most abundant genus (2.5 % average relative abundance across fluid and sediment samples). The site CM2 was dominated by the phylum Bacteroidota and Proteobacteria in higher abundances, with Cyanobacteria, Campylobacterota, and Planctomycetota present at lower abundances. Within Bacteroidota, the most abundant family belonged to *Cyclobacteriaceae*, with no further taxonomic classification (2.3 % average relative abundance across the fluid and sediment samples). Within Proteobacteria, *Woesia* was the most abundant genus (4.3 % average relative abundance across fluid and sediment samples), followed by the SAR 11 and B2M28 clades, with no classification down to the genus level (2.5 % and 2.2 % average relative abundance across the fluid and sediment samples). Within Cyanobacteria, Campylobacterota and Plactomycetota, *Synechococcus* CC9902, *Sulfurovum* and *Rubripirellula* were the most abundant genera (3.5 %, 2.2 %, and 2.2 % average relative abundance across the fluid and sediment samples, respectively).

The BC site located within the Somma-Vesuvius volcanic area, was dominated by members belonging to the Bacteroidota and Proteobacteria phyla, with Desulfurobacterota present at lower abundances (Figure 3). Within Bacteroidota, the most abundant order belongs to the *Flavobacteriales*, with the most abundant ASV not resolved to the genus level (13.3 % average relative abundance across fluid and sediment samples), followed by the clade n7 marine group (9.8 % average relative abundance across fluid and sediment samples), and the genus *Artictiflavibacter* (2.3 % average relative abundance across fluid and sediment samples). The order *Ignavibacteriales* was dominated by the clade LheB3-7 (4.0 % average relative abundance across fluid and sediment samples), followed by the Bacteroidetes VC2.1 Bac22 clade (4.6 % average relative abundance across fluid and sediment samples). Within Proteobacteria, the most abundant genera belong to *Thiomicrorhabdus* (7.8 % average relative abundance across fluid and sediment samples), followed by *Thioclava* (3.8 % average relative abundance across fluid and sediment samples), and *Cocleimonas* (2.7 % average relative abundance across fluid and sediment samples). The low abundance phylum Desulfurobacteriota was dominated by the genus *Desulfobulbus* (2.1 % average relative abundance across fluid and sediment samples).

The biofilm samples were collected in the vicinity of the vent orifice (Figure 4). At the Campi Flegrei region, in SF_G1 and SF_G2 sites (two biofilms were collected - SF_G2A and SF_G2B), the biofilm communities were dominated by ASVs belonging to the Aquificota phyla, more specifically to *Hydrogenothermus* genus (49 %, 37.4 %, and 87.4 % relative abundance, respectively), with other phyla present at lower abundances. On the other hand, at the site CM2, the biofilms were instead dominated by ASVs belonging to the class Gammaproteobacteria (36.9 % relative abundance), with no classification down to the genus level, and the genus *Thiothrix* at lower abundances (8.4 % relative abundance). At the site BC, the most abundant ASVs were affiliated to the genus *Thiothrix* (30.7 % relative abundance).

**Figure 4.**
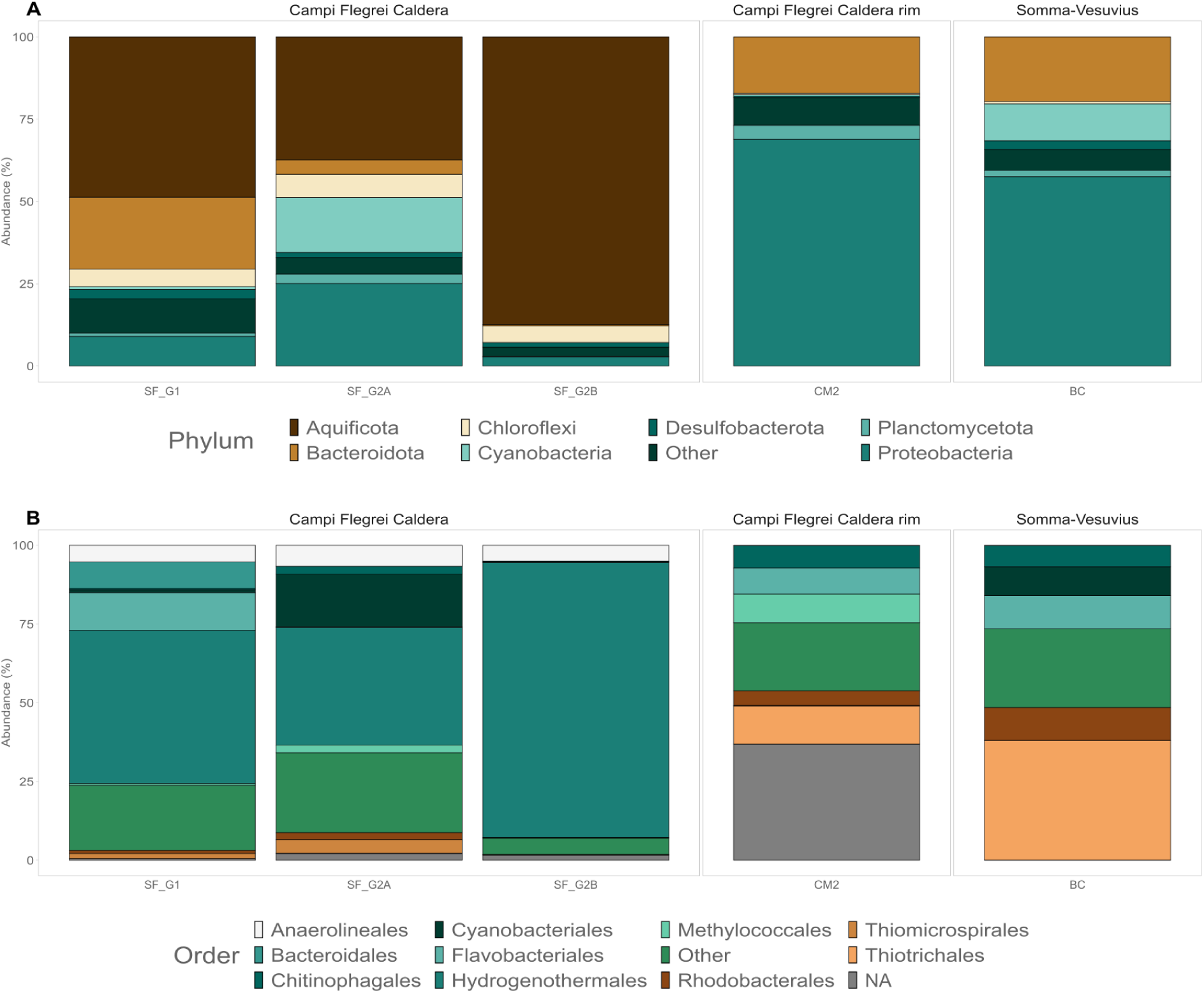
Microbial community composition of the biofilms sampled around the different shallow-water hydrothermal vents studied here: A) Phylum taxonomic rank; B) Order taxonomic rank.

In order to understand the role of the measured geochemical and environmental parameters in constraining the distribution of the microbial communities inhabiting the sampled shallow-water hydrothermal vents, we performed both abundance weighted and unweighted (presence-absence) multivariate ordination (nMDS) based on the Jaccard dissimilarity indexes. We found that the microbial communities are separated according to the different settings where the vents occur, as well as to fluid geochemistry observed in our geochemical analysis (Figure 2A, Figure 5A and 5B, and Supplementary Figure 2 for nMDS based on the unweighted Jaccard index). The clustering based on the geographic location is statistically significant in both the weighted, as well as the unweighted jaccard based nMDS, taking into account the volcanic region (ADONIS, n = 7, p = 0.034 and adj.p = 0.005), as well as the geochemical regime (ADONIS, n = 7, p = 0.034 and adj.p = 0.004). Environmental fitting performed on the weighted Jaccard nMDS show that: salinity, total dissolved solids, conductivity, and the concentrations of Na, Cl, and Br, were statistically correlated with the changes in betadiversity (adj.p < 0.05). Salinity and Sr concentrations were statistically correlated with the unweighted jaccard based nMDS (adj.p < 0.05) (Supplementary Table 2).

**Figure 5.**
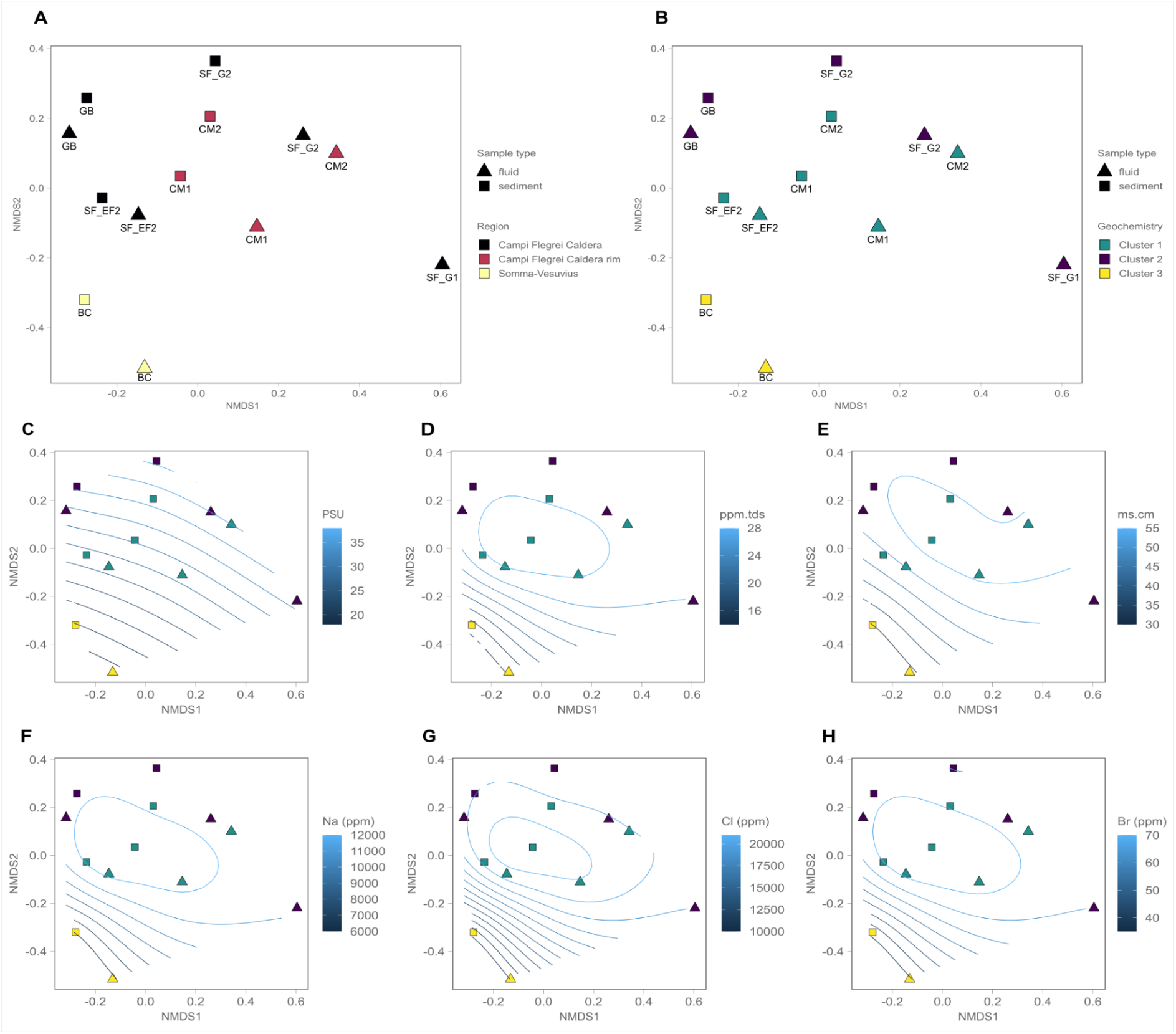
Beta-diversity analysis based on Non-metric multidimensional scaling (nMDS) using the Weighted Jaccard similarity index. A) nMDS coloured by sampling region, B) nMDS coloured by geochemistry (clusters seen on the PCA analysis of the geochemical data) and C-H) *ordisurf* analysis of the most significant variables.

## 4 Discussion

Coupled geochemical and microbiological analyses are fundamental to understanding the geosphere-biosphere feedback in natural ecosystems (Giovannelli et al., 2022). In hydrothermal systems, both continental and marine hosted, geosphere-biosphere coupling is much stronger compared to other ecosystems, since the microbial communities largely rely on compounds and elements provided by the hydrothermal fluids and derived from fluid-rock interactions (Dahle et al., 2018; Fullerton et al., 2021; Barosa et al., 2023; Colman et al., 2023; Basili et al., 2024). In the present study, we analysed the geochemistry of hydrothermal fluids collected from the shallow-water hydrothermal vents of the Gulf of Naples, occurring within two different volcanic regions, Campi Flegrei and Somma-Vesuvius (Figure 1). We also report the geochemical characterization of two new shallow-water hydrothermal vent sites (CM1/2, and GB) that, to our knowledge, have never been described in the literature before. We contextualised the geochemistry of these hydrothermal systems with previous studies done in the region.

### 4.1 Geochemistry of the shallow-water vents of the Gulf of Naples

On the seafloor, the ascent of hydrothermal fluids is facilitated through the permeability of lithospheric fault systems, allowing a more focused channelling of subsurface fluids to the surface (Pearce et al., 2019). All the sampled sites are located within proximity to fault systems (see geological maps in Tramparulo et al., 2018 and Aucelli et al., 2020), which might facilitate both recharging and discharge.

Three major groups of venting fluid geochemistry can be identified using the major ions composition (Table 2 and Figure 2A-E). The first two groups occur within the Campi Flegrei caldera and are hosted in Campania ignimbrite rocks, characterizing the area and can be recognized by their fluid chemistry using classic diagnostic plots for hydrothermal systems (Price et al., 2015). In end-member hydrothermal fluids, Mg^2+^ is depleted due to high temperature water-rock reactions, and it can therefore be used as a tracer to infer the degree of mixing with seawater (Price et al., 2015; Di Napoli et al., 2016). Ion concentrations, including Mg^2+^ in the first group of samples, composed of the fluids from the sites CM1, CM2, and SF_EF, had concentrations and compositions similar to seawater, suggesting a high degree of mixing between the venting hydrothermal fluids and seawater. By contrast, a second group of samples, from GB, SF_G1, and SF_G2, presented a lower degree of mixing, as suggested by the lower Mg^2+^ concentrations and the presence of NH_4_^+^. In seawater, NH_4_^+^ is present in trace amounts, as it is readily assimilated by phytoplankton and oxidised by nitrifying microorganisms (Karl et al., 2008; O’Connor Šraj et al., 2017), while high concentrations of NH_4_^+^ have been reported in hydrothermal springs within the Campi Flegrei region (Sieland, 2009). This suggests a higher contribution of primary hydrothermal fluids to these sites. Additionally, sites SF_G1 and SF_G2 have similar fluid chemistry with respect to previous geochemical data collected in 2006, highlighting the stability of the system (Di Napoli et al., 2016).

The third group, represented only by the site BC, occurs within a different host lithology in the southern portion of the Gulf of Naples (carbonates - Figure 1). The site location in the intertidal zone and the low concentrations of Cl^-^ and Na^+^ compared to the other sites suggest mixing with groundwaters before discharge. A previous study done in the region has shown that the fluids of the vents present in this region are a mixture of seawater and groundwater end-members, as a result of saltwater intrusion into fractured and karstified aquifers, which could account for the differences in fluid chemistry seen here (Baiocchi et al., 2010).

The fluids from the site GB, SF_G1 and SF_G2 were enriched in elements such as Rb, Cs, Fe, As, and Mn. It has been demonstrated that some elements, such as Rb and Cs, can be efficiently extracted from basalt at higher temperatures (Palmer and Edmond, 1989). Cs/Rb ratios in the SF_G1/G2 fluids are high, suggesting equilibration with mafic rocks at depth (Palmer and Edmond, 1989). Sites GB, SF_G1 and SF_G2 were also enriched in As, which can be leached from host lithologies at high temperatures. Arsenic, in the form AsO ^3-^ and AsO ^3-^ is an important electron donor and acceptor for microorganisms and it has already been reported in high concentrations in multiple shallow-water hydrothermal systems (Aiuppa et al., 2006; Pichler et al., 2006; Leal-Acosta et al., 2013; Price et al., 2013; Ruiz-Chancho et al., 2013). Fluids from SF_G1/SF_G2, and GB, were also enriched in Mn and Fe, although with different concentrations (GB site had higher concentrations of Fe and Mn). These elements are frequently enriched in hydrothermal fluids (Tebo and Mandernack, 1993; Scholten et al., 2019), indicating that these fluids could be equilibrating within similar host lithologies. The difference in temperature (more than 30 °C), Fe, and Mn, observed between SF_G1/SF_G2, and GB, could be due to the mixing with seawater as the fluids approach the surface, and/or different water-rock reactions (for instance, in the upflow zone - secondary mineral precipitations), respectively.

The PCA analysis (Figure 2A) shows that the concentrations of the fluids from the shallow-water hydrothermal vents CM1, CM2, and SF_EF cluster together and therefore have similar fluid chemistry. Their similar chemistry, with respect to seawater in the major ions biplots (Figure 2- B-E), suggests a higher degree of mixing at these locations (i.e., a lower ratio between hydrothermal fluid vs seawater). The sites GB, SF_G1, and SF_G2, had similar fluid chemistry between each other (clustering together in the PCA analysis), and were located within the same region within the Campi Flegrei caldera (see Figure 1).

Noble gases, due to their inert properties, can serve as valuable geochemical tracers (Hilton et al., 2002). Helium, for instance, has two stable isotopes: ^3^He and ^4^He. ^3^He is primordial and has been stored in the mantle following Earth’s accretion 4.56 Gyr. In contrast, ^4^He is produced continuously on Earth through the radioactive decay of U and Th (Barry et al., 2022, Hilton et al., 2002). The mean ratio of ^3^He/^4^He for arc gases is approximately 5.4 ± 1.9 Ra (Ra = 1.39 x 10^6^, Hilton et al., 2002; Ozima and Podosek, 2002). Changes in this ratio can provide information about the origin of volatiles in hydrothermal systems (i.e. crust and/or mantle derived) (Hilton et al., 2002). A previous study measured ^3^He/^4^He in both shallow-water hydrothermal vents, as well as in-land geothermal wells and fumaroles in the Campi Flegrei volcanic region (Tedesco et al., 1990). The authors found that there is no significant difference in the ^3^He/^4^He ratio between the sites (^3^He/^4^He ratios ranging between 2.0 and 3.2 Ra), thus suggesting a common source of helium in the two regions. Notably, however, ^3^He/^4^He ratios have been shown to decrease towards the margins of the Campi Flegrei caldera. A lower ^3^He/^4^He ratio in the peripheral manifestations could be due to greater distances from the magmatic source and the primary location of degassing, or mixing with crustal derived volatiles higher in ^4^He (Tedesco et al., 1990). One of the lowest ^3^He/^4^He values measured in Campi Flegrei was found close to where the CM1 and CM2 sites were sampled in the present study. Additionally, Orlando et al. (2011), through the analysis of gas ratios of H_2_, Ar, CH_4_, and CO_2_, found that gases from the Secca della Fumosa region (SF site in this work) are equilibrated at approximately 250 °C, compared to lower equilibration temperatures in samples collected from the Capo Miseno region (150

°C). Even though we did not directly measure gases in this work, the aforementioned studies suggest that the CM1 and CM2 vents occur further away from the main magmatic gas outflow path than, for instance, the vents present in the SF region, supporting regional differences in geochemistry of the fluids controlled by regional difference in volcanism. Since the shape, size and distance to the heat source has been acknowledged to be one of the most important parameters dictating the chemistry of hydrothermal fluids (German and Seyfried, 2014), this could explain both the divergence in fluid chemistry measured in CM1 and CM2, as well as the possible higher mixing with seawater as the fluids approach the surface.

Taken together these results highlight how even in such small scales, regional differences in the host lithologies and geological background can have an impact in the hydrothermal fluid chemistry.

### 4.2 Microbial Diversity

The microbial communities inhabiting the shallow-water hydrothermal vents of the Gulf of Naples, were mostly dominated by ASVs belonging to Proteobacteria and Bacteroidota, with other phyla present at lower abundances (Figure 3). Dominant groups have been previously reported from other shallow-water hydrothermal environments (Zhang et al., 2012; Giovannelli et al., 2013; Gomez-Saez et al., 2017; Price and Giovannelli, 2017, 2019; Barosa et al., 2023). The inherent characteristics of SWHV, such as the presence of both sunlight and geochemical energy, coupled with organic and inorganic carbon sources, reflect both the diversity of taxa as well as the metabolic strategies found in these ecosystems (Price and Giovannelli, 2017). For instance, in the present study, we consistently find ASVs associated with groups known to perform chemolithoautotrophy, chemoheterotrophy, heterotrophy, and photoautotrophy. The differences in their distribution, however, may be related to the topology of the hydrothermal system, as well as to fluid geochemistry and mixing patterns. The mixing of hydrothermal fluids with seawater during ascent and venting creates dynamic mixing zones, generating a plethora of microbial niches with diverse combinations of electron donors and acceptors, suggesting that hydrothermal systems with a higher degree of mixing could support a greater diversity of ecological niches to be explored by the resident microbial communities.

The Campi Flegrei venting sites SF_G1 and SF_G2 were collected at the same location, on two different occasions. They present remarkably different communities. For instance, SF_G1 has the most abundant ASVs related to taxa commonly found in high temperature shallow-water hydrothermal systems (Barosa et al., 2023). At this site, the majority of the ASVs were associated with the genus *Hydrogenothermus*, with the ASV affiliated with the species *Hydrogenothermus marinus* (99.5 % similarity; supplementary table 1). Members belonging to this genus are thermophilic, chemolithoautotrophic bacteria, coupling the oxidation of hydrogen to inorganic carbon fixation using the rTCA cycle (Stöhr et al., 2001). Other abundant ASVs were affiliated with the family *Anaerolineceae*, within Chloroflexi. This family is known to harbour genera employing widely diverse metabolic strategies with anaerobic growth (Bedoya-Urrego and Alzate, 2024). Furthermore, it was also possible to find ASVs related to known groups associated with sulfur metabolism, such as *Thermodesulforhabdus*, which is a thermophilic, acetate-oxidizing, sulfur-reducing bacterium (Beeder et al., 1995), as well as *Thermosulfurimonas,* which is a group known to perform the disproportionation of sulfide to sulfate (Slobodkin et al., 2012). In contrast, samples from site SF_G2 present marked differences in the microbial communities. On one hand, we detected a high abundance of ASVs related to chemolithoautotrophs, such as *Thiomicrospiraceae,* known sulfur-oxidizing thermophilic bacteria, and *Thiogranum,* obligate sulfur-oxidizing, chemolithoautotrophic bacteria (Mori et al., 2015). On the other hand, we found the presence of ASVs affiliated with groups commonly found in open seawater, such as photoautotrophic taxa *Syneccococus* and SAR11 clade, suggesting mixing with seawater. The difference between these two sampling times could be explained by multiple factors, such as changes in the community over time due to the entrainment of seawater in the hydrothermal system or sampling errors that collected a variable amount of seawater during the sampling of SF_G2. The latter seems more plausible given the similar geochemistry and temperature of the fluids (∼68 °C, providing the same selective pressures). Additionally, the microbial communities of the SF_G2 site are more similar to other sampled locations in the study, which are associated with more diffuse and seawater-mixed hydrothermal fluids.

The site SF_EF was sampled approximately 80 meters from SF_G1/2, and was located in an area of vigorous diffuse gas degassing and low temperature anomaly. The geochemistry was very different from the sites SF_G1 and SF_G2, and showed similar chemical composition to seawater (Figure 2-B). At this location, most of the high abundance ASVs are affiliated with known chemoorganotrophs. For instance, the family *Lentimicrobiaceae* has been shown to grow chemoorganotrophically using a narrow range of carbohydrates under anaerobic conditions (Sun et al., 2016). There is a high abundance of ASVs related to known groups that use nitrogen species known to be present in seawater (N_2_O, NO_2_ ^-^, NO ^-^) as electron acceptors. The Bacteroidetes VC2.1Bac22 clade, which is an uncultured clade distributed in marine ecosystems, including hydrothermal systems, has been shown, through MAG analysis, to play key roles in complex organic carbon mineralization. It has also been suggested to play equally important roles in N_2_O sinks in deep-sea hydrothermal systems (Leng et al., 2022, p. 202). High abundance ASVs related to members belonging to the *Calditrix* genus were recovered from the same sample. The type species of this genus, *Calditrix abissi*, couples the oxidation of hydrogen to the reduction of nitrate, using organic carbon to build biomass (Kublanov et al., 2017). The family *Melioribacteraceae* was affiliated with highly abundant ASVs, and are known facultative anaerobic chemoorganotrophic bacteria, which can use Fe^3+^, nitrite (NO_2_ ^-^), and As^5+^, as electron acceptors (Podosokorskaya et al., 2013). The presence of ASVs affiliated with known sulfate-reducing chemoorganotrohic groups was also present, such as *Desulfovibrio*, which are known mesophilic, hydrogenotrophic, sulfate-reducing bacteria previously isolated from a deep-sea hydrothermal system (Alazard et al., 2003). At this site, there were also ASVs affiliated to the species *Methanofolis fontis* (99.47 % similarity; supplementary table 1), which has been isolated from deep-sea sediments, and it is a mesophilic, hydrogenotrophic methanogen (Chen et al., 2020).

Chemolithoautotrophic bacteria are present in high abundances, such as AVS related to the genus *Sulfurovum*, a sulfur and thiosulfate-oxidizing bacterium that uses oxygen or nitrate as the electron acceptor (Inagaki et al., 2004; Giovannelli et al., 2016a). The site GB, from all the sites within the Campi Flegrei caldera, was the closest to land. Similar to the other sites in this region, the ASVs with higher abundances were affiliated with known chemoorganotrophs, chemolithoautotrophs, as well as heterotrophic groups. The most abundant ASVs were affiliated with the species *Marinifilm fragile* (100 % similarity; supplementary Table 1), isolated from a tidal flat sediment in Korea. This bacterium was described as being facultatively anaerobic, moderately halophilic, and was also identified in shallow-water hydrothermal vent systems (Na et al., 2009; Barosa et al., 2023). It was also possible to identify the clade PHOS-HE36, which some previous studies have suggested to be composed of denitrifying bacteria (Xin et al., 2023). A recent study showed that the relative abundance of PHOS-HE36 gradually increased as the Mn^2+^ concentrations increased, as an indication of the strengthening of denitrification (Jiang et al., 2024). This site has the highest concentration of Mn^2+^ measured (Table 3), and therefore indicates a possible relationship between the metals that are being released from the hydrothermal system and the types of metabolism found (Giovannelli, 2023). The family *Cyclobacteriaceae* was also affiliated with abundant ASVs, which have been demonstrated to have wide physiological properties, such as growth temperature and salinity range, and have also been observed in hot springs and mud volcanoes (Dworkin et al., 2006). Similarly to SF_EF, it was possible to identify the family *Melioribacteraceae*. Heterotrophic groups associated with *Thermotomaculum* were affiliated with some ASVs, which are thermophilic, heterotrophic bacteria, initially isolated from deep-sea hydrothermal systems (Izumi et al., 2012). ASVs at this site were also affiliated with the family *Desulfosarcinaceae,* which includes groups associated with sulfate-reducing bacteria (Watanabe et al., 2021). Overall, the shallow-water hydrothermal vents hosted within the Campi Flegrei region were populated by ASVs affiliated with a mixture known chemolithoautotrophic, chemoorganotrophic, and heterotrophic groups, which heavily relied on nitrogen species as electron acceptors, and hydrogen as electron donors, and used organic as well as inorganic carbon to build biomass. These types of metabolisms have been described before in these types of ecosystems in previous studies (Price and Giovannelli, 2017; Barosa et al., 2023).

The shallow-water hydrothermal vents hosted in the Campi Flegrei caldera rim are represented here by the sites CM1 and CM2. These vents have fluid chemistry similar to seawater, suggesting a higher degree of mixing as the fluids approach the surface. At these shallow-water hydrothermal vents, the microbial communities were dominated by ASVs associated with groups known to perform heterotrophic lifestyles, common marine taxa, as well as chemolithoautotrophic and phototrophic groups at lower abundances. In the site CM1, it was possible to identify ASVs affiliated with the *Lutibacter* genus, commonly found in marine environments as well as deep-sea hydrothermal vents, and shown to reduce nitrate to nitrite (Le Moine Bauer et al., 2016). Similar to other shallow-water hydrothermal vents, it was possible to identify ASVs affiliated with *Thiotrichaceae,* which can grow chemolithoautotrophically or chemolithoheterotrophically in the presence of hydrogen sulfide and/or thiosulfate or chemoorganoheterotrophically, as well as *Sulfurovum.* Similarly, at the site CM2, the most abundant ASVs were affiliated with the family *Cyclobacteriaceae*, which are present in a wide variety of environments, including hot springs, and where some species have been shown to degrade polysaccharides and other complex molecules (Dworkin et al., 2006). It was also possible to find ASVs related to common marine heterotrophic bacteria, such as *Woesia*, which have also been recently described in other shallow-water hydrothermal environments (Barosa et al., 2023). At this site, we also retrieved ASVs related to known photoautotrophic taxa, *Synechococcus* CC9902, that is widely distributed in marine environments, and is regarded as one of the most important components of photosynthetic picoplankton (Kim et al., 2018), as well as SAR11, chemoheterotrophic alphaproteobacteria accounting for 25 % of all plankton (Giovannoni, 2017). Notwithstanding, there were still present ASVs related to chemolithoautotrophic taxa, such as in the case of *Sulfurovum.* At both these sites, the increase of groups associated with heterotrophic metabolisms in the high abundance ASVs, as well as common marine taxa, sheds light on the complexities inherent to the mixing ratios observed in shallow-water hydrothermal systems. Vents hosted further away from a heat source, and where hydrothermal fluids mix to a higher degree with seawater, present lower temperatures and the presence of increased members associated with marine taxa and heterotrophic metabolisms. This pattern was also observed in the Aeolian archipelago, especially in the islands with less pronounced volcanic activity (Barosa et al., 2023).

The site BC was located within the area of influence of the Somma-Vesuvius volcano. The geochemistry of these fluids was the most divergent compared to all the other sites of the Campi Flegrei region. As discussed in the geochemistry section, these differences could be due to difference in the host rocks between BC, occurring in carbonates in contact with volcanic deposits, compared to the Campi Flegrei sites occurring within trachytic ignimbrites, by diverse primary fluids and mixing regimes between the two areas, or most likely a combination of both. In the BC site was possible to find the presence of ASVs related to common marine taxa, as well as high abundances of ASVs affiliated to groups capable of sulfur oxidation, such as the genus *Thiomicrorhabdus, Thioclava,* and *Cocleimonas* (Sorokin et al., 2005; Tanaka et al., 2011; Kojima and Fukui, 2019). From all the sites, BC is the one with the lowest oxidation reduction potential, suggestive of a highly reducing environment. This is also supported by the very low concentrations of SO_4_ ^2-^, coupled with a very strong smell of “rotten eggs” at the location, characteristic of the presence of H_2_S. Enzymes catalyzing the oxidation of sulfur species, as noted in (Hay Mele et al., 2023), are heavily based on the trace metals Fe and Mo. These specific metals were present in very low abundances at this site, especially Mo, below the detection limit in all dilutions conducted pre-measurements. Microbial communities inhabiting strongly reducing environments undergo selective pressures to exploit the abundance of “free” electrons for their metabolic needs. This could imply heavy scavenging of the specific metals needed to perform these reactions, possibly explaining their low abundance in this particular site. The relationship between metal availability and microbial distribution and metabolism has been suggested in recent studies, and this could provide yet another hint on how these relationships happen in complex natural environments (Giovannelli, 2023; Hay Mele et al., 2023).

At sites SF_G1 and SF_G2 in the Campi Flegrei region, the biofilms sampled in the vicinity of the vents were mostly composed of ASVs affiliated with the Aquificota phylum, more specifically with the genus *Hydrogenothermus.* Interestingly, previous studies that analyzed the microbial communities of biofilms in both deep-sea and shallow-water hydrothermal systems report a community composed mainly of sulfur-oxidizing groups belonging to Campylobacterota and the class Gammaproteobacteria (Gulmann et al., 2015; O’Brien et al., 2015; Sciutteri et al., 2022; Barosa et al., 2023). Members belonging to the Aquificota, on the other hand, have been identified in biofilms of high temperature deep-sea vents (Fullerton et al., 2024). This suggests that both the availability of electron donors and acceptors, as well as the temperature of the system, can have important roles in constraining the establishment of bacterial biofilms in natural hydrothermal systems. For this reason, Secca delle Fumose, due to its easy access, can serve as an ideal laboratory to understand biofilm dynamics in hydrothermal systems.

The beta-diversity analysis (Figure 5) coupled with permutational multivariate analysis (ADONIS) shows that the microbial communities from the shallow-water hydrothermal vents hosted in different settings, that is, the Campi Flegrei caldera, Campi Flegrei caldera rim, and the Somma-Vesuvius regions, are statistically different from one another (Figure 5A). That takes into account the most abundant groups, as well as the rare species. Interestingly, the vents separated by type of fluid chemistry are also statistically different. Differences between microbial communities inhabiting geologically and geographically distinct shallow-water hydrothermal vents have been previously reported for the shallow vents of the Aeolian archipelago (Barosa et al., 2023). This suggests that the geological setting where the vents occur and the consequent difference in hydrothermal fluid chemistry can impose controls on microbial distribution. Geological controls on the microbial diversity have also been observed on land for deeply sourced seeps in the volcanic regions of Costa Rica, Peru, and Bolivia (Fullerton et al., 2021; Colman et al., 2023; Sims et al., 2023; Upin et al., 2023). In SWHV, however, the complex mixing with seawater, the presence of sunlight, tidal forcing, and organic matter inputs to the system, could overlap, cross and influence geological and geochemical processes, also playing a key role in mediating microbial diversity and distribution. These parameters are in themselves a *property* of each individual hydrothermal system. For instance, recent studies have demonstrated that in deeply sourced seeps of Costa Rica, even though chemolithoautotrophic communities were responding to changes in geochemistry, no such pattern was observed for secondary consumers, i.e, heterotrophs (Paul et al., 2023). The quality and abundance of fresh organic matter might play an important role in controlling the abundance of taxa associated with heterotrophic metabolic strategies. In SWHV, organic matter can be either produced by chemolithoautotrophs, phototrophs, as well as imported to the system from the water column or terrigenous input from nearby landmasses (Price and Giovannelli, 2017). In the shallow-water hydrothermal vents of the Aeolian archipelago, the presence of fresh organic matter in some sites was associated with the higher abundance of heterotrophic groups (Barosa et al., 2023). Gases from volcanic origin, such as H_2_, CO_2_, H_2_S, and CH_4_, due to their key roles in microbial redox metabolism as electron donors and acceptors, might also play a key role in shaping ecological niches for microbial exploitation in these systems.

The *ordisurf* analysis conducted here (Figure 5C-H) shows how the groups that have similar fluid chemistry (as seen in the PCA analysis on the geochemistry section), cluster closely, illustrating their similar microbial community structure. As previously discussed, it follows that the nature of any given hydrothermal system, such as its proximity to the magmatic heat source, has a direct influence on the types of temperature dependent water-rock-gas reactions happening during fluid trajectory, and consequent fluid chemistry, dictating also the degree to which the upflowing fluids mix with seawater. This highlights the importance of taking a multidisciplinary sampling approach to study geothermal environments, such as the one suggested by Giovannelli at al., 2022. Additionally, in order to untangle the complex mixing ratios in these systems, we further suggest an approach based on the background concept introduced in Cascone at al., 2025. By sampling background seawater, and similar to what is currently done in geochemical analysis, it can allow us to provide a microbiological “end-member” to gain further knowledge on life inhabiting these dynamic ecosystems.

## 5 Conclusion

The Gulf of Naples shows intense underwater hydrothermalism as a function of the volcanic activity in the area. In the present study, we describe both the geochemistry as well as the microbiology of the SWHV occurring within two different volcanic regions, Campi Flegrei and Somma-Vesuvius. Through geochemical analysis, we found that on the SWHV sampled, hydrothermal fluid chemistry is contingent on the region where the vents occur, with the distance from the heat source possibly playing a key role in dictating the chemistry of the fluids, corroborated by gas geochemistry measurements conducted in the past. Additionally, we found diverse microbial consortia inhabiting these SWHV, with a high abundance of ASVs affiliated with known chemolithoautotrophic, heterotrophic, and photoautotrophic groups. This is consistent with previous studies conducted in other shallow-water hydrothermal systems, and therefore, can be regarded as a definite characteristic of these dynamic systems. Notably, here the distribution of the ASVs is suggested to be related to differences in hydrothermal fluid geochemistry as a function of fluid mixing landscapes, as well as the vent’s location (Campi Flegrei caldera, Campi Flegrei caldera rim, and Somma-Vesuvius). The chemistry of hydrothermal fluids, as a response to water-rock-gas reactions at depth, can strongly influence the availability/diversity of ecological niches to be exploited by microorganisms, through differences in electron donors and acceptors, as well as organic and inorganic carbon sources. On the other hand, the geological background and location of hydrothermal systems, and the consequent diffusivity of the fluids approaching the surface, can generate complex mixing ratios with seawater, which, coupled with sunlight, tidal fluctuations, and organic matters, can also play key roles in constraining microbial distribution and composition in SWHV. As the Campi Flegrei volcano continues to exhibit signs of unrest with increasing seismic activity, this work provides an important baseline for future studies aiming at understanding how microbial communities respond to geochemical shifts driven by volcanic processes. In this context, our findings could contribute valuable insights for the development of long-term monitoring strategies within highly geodynamic systems.

## 6 Data Availability Statement

Sequences are available through the European Nucleotide Archive (ENA) with bioproject accession number PRJEB67762 under the Umbrella Project CoEvolve PRJEB55081. A complete R script containing all the steps to reproduce our analysis is available at https://github.com/giovannellilab/Barosa_et_al_FEAMP_shallow_vent_diversity permanently stored with DOI: 10.5281/zenodo.14966493.

## 8 Author Contributions

BB carried out data analysis, geochemical experiments and wrote the first draft of the manuscript. BB, CC, MS, SD, MC, AB, MCO, and LD produced the geochemical and microbiological data. CC performed DNA extractions. CC, MS, DB, MC, RP, CV, and DG planned the expedition and collected the samples. RP, CV, AC, and DG supervised the study. All authors contributed equally to the final version of the manuscript.

## Acknowledgments

The authors wish to thank the Pozzuoli diving centre (Centro Sub Pozzuoli) and Guglielmo Fragale for providing diving logistics necessary to carry out the sampling.

