## Supplementary material for "The microbiology and geochemistry of the shallow-water hydrothermal vents of the Gulf of Naples, Italy"

**Supplementary figure 1-** Correlation plot between all the variables measured in the present study

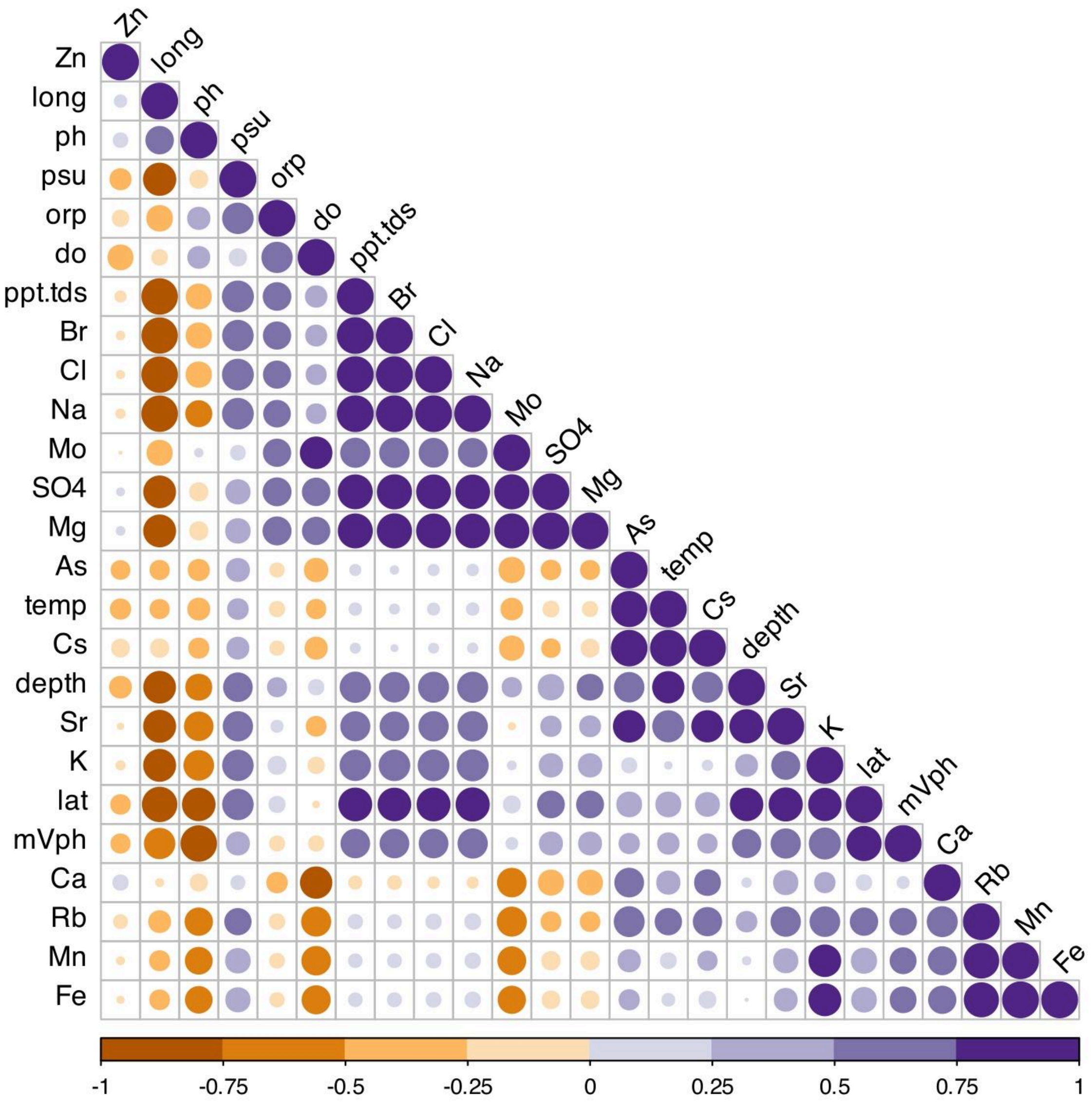

**Supplementary figure 2-** Beta-diversity analysis based on Non-metric multidimensional scaling (nMDS) using the Unweighted Jaccard similarity index colored by 1) Region and 2) Geochemistry of the fluids

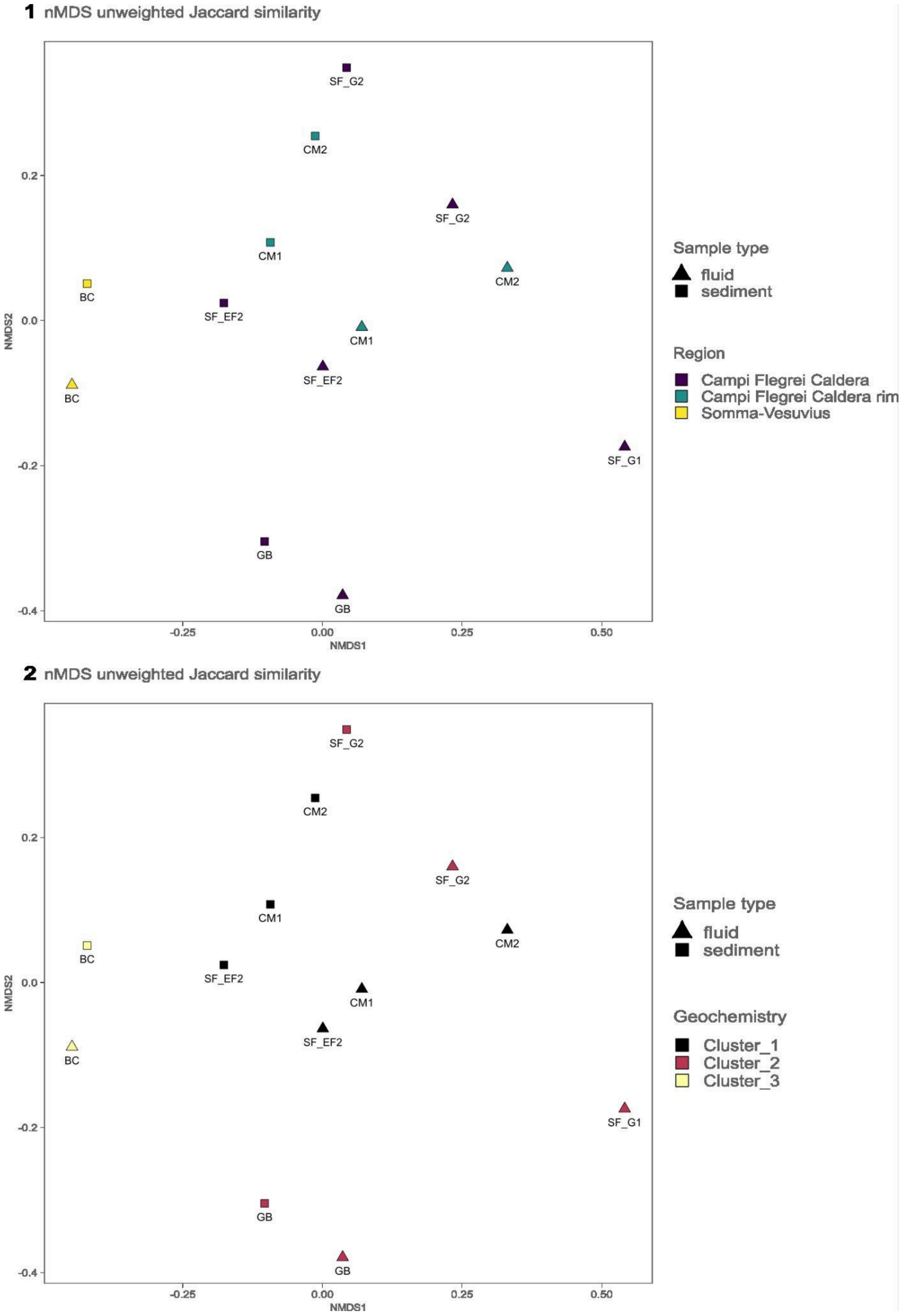

**Supplementary table 1**. Table of the major ASVs (above 2 %) at each of the sampling sites (Abundance % is reported as an average between fluid and sediment samples. Taxonomic classifications of Silva, Ez taxon, and NCBI nucleotide database.

| ASV | Code | Abundance (%) | Silva classification | Ez taxon classification | Similarity (%) | Accession | NCBI nucleotide classification | Similarity (%) | Accession | Environment |
|---|---|---|---|---|---|---|---|---|---|---|
| TACGGAGGGTGCAAGCGTTATCCGGATTCATTGGGTTTAAAGGGTGCGCAGGCGGACTTTTAAGTCAGTGGTGAAATCCCGGGGCTCAACCCCGGAACTGCCATTGATACTGAAAGTCTTGAGTTTGGTTGAAGTAGGCGGAATGTAGCATGTAGCGGTGAAATGCTTAGATATGCTACAGAACACCGATCGCGAAGGCAGCTTACTAAACCAATACTGACGCTCAGGCACGAAAGCGTGGGGAGCGAACAGGATTAGATACCCTGGTAGTCCACGCCGTAAACTATGATCACTCGTTGTTGGCGATACACAGTCAGCGACCAAGCGAAAGCGATAAGTGATCCACCTGGGGAGTACGATCGCAAGGTTG | BC | 4..58 | Bacteroidetes_VC2.1_Bac22 | AY197392_s | 98.87 | AY197392 | Uncultured Cytophaga sp. | 100.00 | AB238986.1 | deep-sea cold seep sediments |
| TACGGAGGGTGTAAGCGTTATCCGGAATCATTGGGTTTAAAGGGTCTGTAGGCGGATTGCTAAGTCAGGGGTGAAATCCCACAGCTCAACTGTGGCATTGCCTTTGATACTGGTGATCTTGAGATATAGTGAGGTAGATAGAATGTGTAGTGTAGCGGTGAAATGCATAGATATTACACAGAATACCGATTGCGAAGGCAGTCTACTAACTATCATCTGACGCTGATAGACGAAAGCGTGGGGAGCGAACAGGATTAGATACCCTGGTAGTCCACGCCGTAAACGATGGATACTAGCTGTTGGACTTTAGGGTTCAGTGGCCAAGCGAAAGTGATAAGTATCCCACCTGGGGAGTACGTTCGCAAGAATG | BC | 13.13 | *Flavobacteriaceae* | EU487997_s | 98.65 | EU487997 | Uncultured bacterium clone CK_1C3_19 | 98.6 | EU487997.1 | siliciclastic sediment from Thalassia seagrass bed |
| | BC | 9.45 | NS7_marine_group | FQ032827_s | 93.78 | FQ032827 | Uncultured marine | 95.14 | KM223594.1 | Ultra-oligotrophic |

| | | | | | | | | | | |
|---|---|---|---|---|---|---|---|---|---|---|
| TACGGAGGGTGCAAGCGTTATCCGGAATCATTGGGTTTAAAGGGTGCGTAGGCGGAAATTTAAGTCAGTGGTGAAAGCCCACAGCTCAACTGTGGAACTGCCATTGATACTGAATTTCTTGAATATGATTGAAGTAGGCGGAATGTGTCATGTAGCGGTGAAATGCATAGATATGACACAGAACACCGATAGCGAAGGCAGCTTACTAAGTCATTATTGACGCTGAGGCACGAAAGCGTGGGGAGCGAACAGGATTAGATACCCTGGTAGTCCACGCCGTAAACTATGATTACTCGTTCTTGGCGATACACAGTCAGGGACTAAGCGAAAGTGATAAGTAATCCACCTGGGGAGTACGATCGCAAGGTTG | | | | | | | bacterium clone BIOS04_MAR_15m171 | | | South Pacific Ocean |
| TACAGAGGGTGCAAGCGTTATTCGGAATTACTGGGCGTAAAGCGCGCGTAGGCGGATTATTAAGTCAGTTGTGAAAGCCCTGGGCTCAACCTGGGAACTGCATCTGATACTGGTAGTCTAGAGTTTAAGAGAGGGAAGTGGAATTCCAGGTGTAGCAGTGAAATGCGTAGATATCTGGAGGAACATCAGTGGCGAAGGCGACTTCCTGGCTTAAAACTGACGCTGAGGTGCGAAAGCGTGGGTAGCGAACGGGATTAGATACCCCGGTAGTCCACGCCGTAAACGATGTCAACTAGTTGTTGGTCTTATTAAAAAGATTAGTAACGAAGCTAACGCGATAAGTTGACCGCCTGGGGAGTACGGTCGCAAGATTA | BC | 7.83 | *Thiomicrorhabdus* | *Thiomicrorhabdus hydrogeniphila* | 100 | LC010781 | Uncultured bacterium clone D1-STK-23 | 98.37 | NR_028679.1 | NA |
| TACGTAGGGGGCAAGCGTTGTCCGGATTTACTGGGTGTAAAGGGCGCGTAGGCGGGTTTGTAAGTCAGTGGTGAAATCCTACAGCTTAACTGTAGAACTGCCTTTGATACTGCAGACCTTGAGTACGGAAGAGAGAGGCGGAATTCCAGGTGTAGTGGTGAAATACGTAGATATCTGGAAGAACACCAGAGGCGAAGGCGGTCTCTTGGTCCGTTACTGACGCTGAGGCGCGAAAGCGTGGGGAGCAAACAGGATTAGATACCCTGGTAGTCCACGCCGTAAACGATGAATACTAGGTGTTGGGTTTTTAACTCAGTGCCGCAGCAAACGCATTAAGTATTCCACCTGGGGAGTACGATCGCAAGGTTG | BC | 4.01 | IheB3-7 | JQ579954_s | 97.83 | JQ579954 | *Melioribacter roseus* P3M-2 | 92.49 | NR_074796.1 | surface of a chute, under the flow of hot water coming from an oil exploring well |

| | | | | | | | | | | |
|---|---|---|---|---|---|---|---|---|---|---|
| TACGGAGGGGGCTAGCGTTGTTCGGAATTACTGGGCGTAAAGCGCACGTAGGCGGATTATTAAGTGAGGGGTGAAATCCCAGGGCTCAACCCTGGAACTGCCTCTCATACTGGTAGTCTTGAGTTCGAGAGAGGTGAGTGGAATTCCGAGTGTAGAGGTGAAATTCGTAGATATTCGGAGGAACACCAGTGGCGAAGGCGGCTCACTGGCTCGATACTGACGCTGAGGTGCGAAAGCGTGGGGAGCAAACAGGATTAGATACCCTGGTAGTCCACGCCGTAAACGATGAATGCCAGTCGTCGGGCAGTATACTGTTCGGTGACACACCTAACGGATTAAGCATTCCGCCTGGGGAGTACGGTCGCAAGATTA | BC | 3.8 | *Thioclava* | *Thioclava indica* | 98.92 | AUNB01000081 | *Thioclava indica* strain MCCC 1A00513 | 98.92 | NR_136454.1 | surface seawater of the Indian Ocean |
| TACGGAGGATCCAAGCGTTATCCGGAATCATTGGGTTTAAAGGGTCCGTAGGTTGATAATTAAGTCAGAGGTGAAATCCTGCCGCTCAACGGTAGAATTGCCTTTGATACTGGTTATCTTGAGTTATTATGAAGTAGTTAGAATATGTAGTGTAGCGGTGAAATGCATAGATATTACATAGAATACCAATTGCGAAGGCAGATTACTAATAATGAACTGACACTGATGGACGAAAGCGTGGGGAGCGAACAGGATTAGATACCCTGGTAGTCCACGCCGTAAACGATGGATACTAGCTGTTCGGATTTCGGTCTGAGTGGCTAAGCGAAAGTGATAAGTATCCCACCTGGGGAGTACGTTCGCAAGAATG | BC | 2.76 | *Arcticiflavibacter* | *Flavivirga amylovorans* | 98.1 | HM475138 | *Wocania arenilitoris* strain HMF6543 | 97.84 | NR_159135.1 | Seashore Sand |
| TACAGAGGGTGCAAGCGTTAATCGGAATTACTGGGCGTAAAGCGCGCGTAGGCGGTTAGATAAGTCAGATGTGAAATCCCCGGGCTCAACCTGGGAACTGCACCTGATACTGTCTGGCTAGAGTTTTGGAGAGGGAAGTAGAATTTCAGGTGTAGCGGTGAAATGCATAGAGATCTGAAGGAATACCAGTGGCGAAGGCGACTTCCTGGCCAAAAACTGACGCTGAGGTGCGAAAGCGTGGGTAGCAAACGGGATTAGATACCCCGGTAGTCCACGCCGTAAACGATGTCAACTAGTCGTTGGTTCCCTTGAGGGATCGGTGACGCAGCTAACGCATTAAGTTGACCGCCTGGGGAGTACGGTCGCAAGACTA | BC | 2.73 | *Cocleimonas* | *Cocleimonas flava* | 98.12 | AB495251 | *Cocleimonas flava* strain KMM 3898 | 98.12 | NR_112909.1 | sand snail |

| | | | | | | | | | | |
|---|---|---|---|---|---|---|---|---|---|---|
| TACGGAGGGTGCAAGCGTTACT<br>CGGAATCACTGGGCGTAAAGCG<br>CATGCAGGCGGTTTAATAAGTTA<br>GAAGTGAAATCCTACAGCTTAA<br>CTGTAGAACTGCTTCTAAAACTG<br>TTAGACTAGAGTCTGGGAGGGG<br>AAGATGGAATTAGTAGTGTAGG<br>GGTAAAATCCGTAGAGATTACTA<br>GGAATACCAAAAGCGAAGGCGA<br>TCTTCTGGAACAGTACTGACGCT<br>GAGATGCGAAAGCGTGGGGAGC<br>AAACAGGATTAGATACCCTGGTA<br>GTCCACGCAGTAAACGATGAAT<br>GTTAGTCGTCGGGGCACTAGTT<br>GTTTCGGTGATGCAGTTAACACA<br>TTAAACATTCCGCCTGGGGAGTA<br>CGGTCGCAAGATTA | BC | 2.3 | *Sulfurovum* | Maps-EB01 | 96.51 | AB278150 | *Sulfurovum denitrificans strain eps51* | 92.80 | NR_179236.1 | hydrothermal field |
| TACGGAGGGTGCAAGCGTTGTT<br>CGGAATCACTGGGCGTAAAGGG<br>CGCGCAGGCGGTTTGATTAGTC<br>AGATGTGAAAGCCCACGGCTTA<br>ACCGTGGAAGTGCATTTGAAAC<br>TGTCAGACTTGAGTATCAGAGG<br>GGAAAGTGGAATTCCCGGTGTA<br>GAGGTGAAATTCGTAGATATCGG<br>GAGGAATACCGGTGGCGAAGGC<br>GACTTTCTGGCTGAATACTGACG<br>CTGAGGCGCGAAAGCGTGGGG<br>AGCAAACAGGATTAGATACCCT<br>GGTAGTCCACGCCGTAAACGAT<br>GTCAACTAGGTGTAGGGGGTGT<br>TGATCCCCTCTGTGCCGCAGCTA<br>ACGCATTAAGTTGACCGCCTGG<br>GGAGTACGGTCGCAAGATTA | BC | 2.13 | *Desulfobulbus* | AF449227_s | 99.73 | AF449227 | *Desulfobulbus elongatus* strain FP | 97.86 | NR_029305.1 | NA |
| TACGGAGGGTGCAAGCGTTATC<br>CGGAATCATTGGGTTTAAAGGGT<br>CCGTAGGCGGATTGATAAGTCA<br>GAGGTGAAATCCCACAGCTTAA<br>CTGTGGCACTGCCTTTGATACTG<br>TTAGTCTTGAATTATATGGAAGT<br>AGATAGAATGTGTAGTGTAGCGG<br>TGAAATGCTTAGAGATTACACAG<br>AATACCGATTGCGAAGGCAGTC<br>TACTACGTATATATTGACGCTAAT<br>GGACGAAAGCGTGGGGAGCGA<br>ACAGGATTAGATACCCTGGTAGT<br>CCACGCCGTAAACGATGGACAC<br>TAGTTGTTGGATTTATTCAGTGA<br>CTAAGCGAAAGTGATAAGTGTC<br>CCACCTGGGGAGTACGATCGCA<br>AGATTG | CM1 | 7.98 | *Lutibacter* | *Lutibacter holmesii* | 97.81 | JQ241142 | Uncultured bacterium clone MethaneSIP16-4-44 | 99.73 | GU584454.1 | Marine Hydrocarbon Seeps |

| | | | | | | | | | | |
|---|---|---|---|---|---|---|---|---|---|---|
| TACGGAGGGTGTAAGCGTTATCC<br>GGAATCATTGGGTTTAAAGGGTC<br>TGTAGGCGGATTGCTAAGTCAG<br>GGGTGAAATCCCACAGCTCAAC<br>TGTGGCATTGCCTTTGATACTGG<br>TGATCTTGAGATATAGTGAGGTA<br>GATAGAATGTGTAGTGTAGCGGT<br>GAAATGCATAGATATTACACAGA<br>ATACCGATTGCGAAGGCAGTCTA<br>CTAACTATCATCTGACGCTGATA<br>GACGAAAGCGTGGGGAGCGAA<br>CAGGATTAGATACCCTGGTAGTC<br>CACGCCGTAAACGATGGATACTA<br>GCTGTTGGACTTTAGGGTTCAGT<br>GGCCAAGCGAAAGTGATAAGTA<br>TCCCACCTGGGGAGTACGTTCG<br>CAAGAATG | CM1 | 5.14 | *Flavobacteriaceae* | EU487997_s | 98.65 | EU487997 | Uncultured bacterium clone CK_1C3_19 | 98.65 | EU487997.1 | geochemical habitat on bacterial symbiont |
| TACGGAGGGTGCAAGCGTTATC<br>CGGAATCACTGGGTTTAAAGGG<br>TGCGTAGGTTGTTAGATAAGTCA<br>GAGGTGAAAGCCTGCAGCTAAA<br>CTGTAGAACTGCCTTTGATACTG<br>TCTGGCTTGAATTAGGTTGAGGT<br>TGGCGGAATGTGACATGTAGCG<br>GTGAAATGCATAGATATGTCATA<br>GAACACCAAATGCGAAGGCAGC<br>TGACTAGACCTTAATTGACACTG<br>AGGCACGAAAGCGTGGGGAGC<br>GAACAGGATTAGATACCCTGGTA<br>GTCCACGCCCTAAACGATGCTTA<br>CTGGATGTGTGCCCTTCGGGGT<br>GCGCATCTGAGGGAAACCATTA<br>AGTAAGCCACCTGGGGAGTACG<br>TCGGCAACGATG | CM1 | 4.21 | *Saprospiraceae* | JQ579834_s | 92.95 | JQ579834 | Uncultured Bacteroidetes bacterium clone FII-TR039 | 92.70 | JQ579834.1 | oil-polluted subtidal sediments |
| TACGGAGGGGGTTAGCGTTGTT<br>CGGAATTACTGGGCGTAAAGCG<br>CACGTAGGCGGACTGGAAAGTT<br>GGGGGTGAAATCCCGGGGCTCA<br>ACCCCGGAACTGCCTCCAAAAC<br>TTCCAATCTAGAGATCGAGAGA<br>GGTGAGTGGAATTCCAAGTGTA<br>GAGGTGAAATTCGTAGATATTTG<br>GAGGAACACCAGTGGCGAAGG<br>CGGCTCACTGGCTCGATACTGAC<br>GCTGAGGTGCGAAAGCGTGGGG<br>AGCAAACAGGATTAGATACCCT<br>GGTAGTCCACGCCGTAAACGAT<br>GAGAGCTAGACGTCGGGCAGCA | CM1 | 3.84 | Rhodobacteraceae | EU491911_s | 99.46 | EU491911 | Uncultured bacterium clone G46 | 100.00 | JX568047.1 | |

| | | | | | | | | | | |
|---|---|---|---|---|---|---|---|---|---|---|
| TGCTGTTCGGTGTCGCAGTTAAC GCATTAAGCTCTCCGCCTGGGG AGTACGGTCGCAAGATTA | | | | | | | | | | |
| TACGGAGGATCCAAGCGTTATCC GGAATCATTGGGTTTAAAGGGTC CGTAGGTGGATAATTAAGTCAGA GGTGAAATCCTGCAGCTCAACT GTAGAATTGCCTTTGAAACTGGT TATCTTGAGTTATTATGAAGTAGT TAGAATATGTAGTGTAGCGGTGA AATGCATAGATATTACATAGAATA CCAATTGCGAAGGCAGATTACTA ATAATATACTGACACTGATGGAC GAAAGCGTGGGGAGCGAACAG GATTAGATACCCTGGTAGTCCAC GCCGTAAACGATGGTCACTAGC TGTTCGAACTTCGGTTTGAGTG GCTAAGCGAAAGTGATAAGTGA CCCACCTGGGGAGTACGTTCGC AAGAATG | CM1 | 3.67 | *Olleya* | *Olleya namhaensis* | 99.73 | Jgi.1108060 | *Olleya* sp. R2A056317 | 99.73 | LR722880.1 | Coastal marine surface water |
| TACGGAGGGTGCAAGCGTTAAT CGGAATTACTGGGCGTAAAGGG TACGTAGGCGGTTATTTAAGTCA GATGTGAAATCCCTGGGCTCAA CCTAGGAATGGCATCTGATACTG GATAACTTGAGTTTAAGAGAGG AGTGTGGAATTTCCGGTGTAGC GGTGAAATGCATAGAGATCGGA AGGAACATCAGTGGCGAAGGCG GCACTCTGGCTTAAAACTGACG CTGAGGTACGAAAGCGTGGGTA GCAAACAGGATTAGATACCCTG GTAGTCCACGCCCTAAACGATGT CAACTAGCCGTTGGATCCATTTA AGGATTTAGTGGTGCAGCTAAC GCATTAAGTTGACCGCCTGGGG AGTACGCACGCAAGTGTA | CM1 | 3 | *Thiotrichaceae* | GU369918_s | 99.2 | GU369918 | Uncultured gamma proteobacterium clone V1B07b16 | 99.20 | GU369918.1 | shallow hydrothermal Vents |

| | | | | | | | | | |
|---|---|---|---|---|---|---|---|---|---|
| TACGGAGGGTCCAAGCGTTATC<br>CGGATTTATTGGGTTTAAAGGGT<br>CCGTAGGCGGGGTTTTAAGTCA<br>GTGGTGAAATCCTACAGCTCAA<br>CTGTAGAACTGCCATTGAAACT<br>GGAACTCTTGAATGTGATTGAG<br>GTAGGCGGAATATGTCATGTAGC<br>GGTGAAATGCTTAGATATGCCAT<br>AGAACACCGATAGCGAAGGCAG<br>CTTACCAAGTCATAATTGACGCT<br>GATGGACGAAAGCGTGGGGAGC<br>GAACAGGATTAGATACCCTGGTA<br>GTCCACGCCGTAAACGATGATC<br>ACTAGCTATTGGCGATATACAGT<br>CAGTGGCACAGCGAAAGTGTTA<br>AGTGATCCACCTGGGGAGTACG<br>ATCGCAAGGTTG | CM1 | 2.6 | *Cryomorphaceae* | AY225660_s | 95.95 | AY225660 | Uncultured bacterium NZ-BA-8 | 98.11 | AB239762.1 | Brothers Caldera, south Kermadec Arc |
| TACGGAGGGTGCAAGCGTTACT<br>CGGAATCACTGGGCGTAAAGCG<br>CATGCAGGCGGTTTAATAAGTTA<br>GAAGTGAAATCCTACAGCTTAA<br>CTGTAGAACTGCTTCTAAAACTG<br>TTAGACTAGAGTCTGGGAGGGG<br>AAGATGGAATTAGTAGTGTAGG<br>GGTAAAATCCGTAGAGATTACTA<br>GGAATACCAAAAGCGAAGGCGA<br>TCTTCTGGAACAGTACTGACGCT<br>GAGATGCGAAAGCGTGGGGAGC<br>AAACAGGATTAGATACCCTGGTA<br>GTCCACGCAGTAAACGATGAAT<br>GTTAGTCGTCGGGGCACTAGTT<br>GTTTCGGTGATGCAGTTAACACA<br>TTAAACATTCCGCCTGGGGAGTA<br>CGGTCGCAAGATTA | CM1 | 2.5 | *Sulfurovum* | AB278150_s | 96.51 | AB278150 | Uncultured bacterium clone G56 | 98.39 | JX568056.1 | marine hydrocarbon seep |
| TACGGAGGGGGTTAGCGTTGTT<br>CGGAATTACTGGGCGTAAAGCG<br>CACGTAGGCGGATTAGTCAGTC<br>AGAGGTGAAATCCCAGGGCTCA<br>ACCCTGGAACTGCCTTTGATACT<br>GCTAGTCTTGAGTTCGAGAGAG<br>GTGAGTGGAATTCCGAGTGTAG<br>AGGTGAAATTCGTAGATATTCGG<br>AGGAACACCAGTGGCGAAGGC<br>GGCTCACTGGCTCGATACTGAC<br>GCTGAGGTGCGAAAGCGTGGGG<br>AGCAAACAGGATTAGATACCCT<br>GGTAGTCCACGCCGTAAACGAT<br>GAATGCCAGACGTCGGGTAGCA<br>TGCTATTCGGTGTCACACCTAAC<br>GGATTAAGCATTCCGCCTGGGG<br>AGTACGGTCGCAAGATTA | CM1 | 2.1 | *Actibacterium* | *Aliiruegeria sabulilitoris* | 98.92 | LOAS01000080 | Uncultured bacterium clone SAPTA-27 | 100.00 | MG251626.1 | tropical mangrove |

| | | | | | | | | | | |
|---|---|---|---|---|---|---|---|---|---|---|
| TACGGAGGGTGCAAGCGTTATC<br>CGGAATCATTGGGTTTAAAGGGT<br>CCGTAGGCGGGTCATTAAGTCA<br>GAGGTGAAATCCCACAGCTTAA<br>CTGTGGAACTGCCTTAGATACTG<br>ATGATCTTGAGTTTTAGTGAAGT<br>AGATAGAATGTGTAGTGTAGCGG<br>TGAAATGCATAGATATTACACAG<br>AATACCGATTGCGAAGGCAGTC<br>TACTAACTAACAACTGACGCTAA<br>TGGACGAAAGCGTGGGGAGCG<br>AACAGGATTAGATACCCTGGTAG<br>TCCACGCCGTAAACGATGGATAC<br>TAGTTGTTTGAGATTTATCTTGA<br>GTGACTAAGCGAAAGTGATAAG<br>TATCCCACCTGGGGAGTACGTTC<br>GCAAGAATG | CM1 | 2 | *Maritimimonas* | *JQ579897_s* | 100 | JQ579897 | Uncultured Bacteroidetes bacterium clone FII-TR102 | 100.00 | JQ579897.1 | oil-polluted subtidal sediments |
| TACGGAGGGTGCAAGCGTTAAT<br>CGGAATTACTGGGCGTAAAGCG<br>CGCGTAGGCGGCTTTGTAAGTC<br>GGATGTGAAATCCCCGGGCTCA<br>ACCCGGGAACTGCATTCGATACT<br>GCAGAACTAGAGTATGGTAGAG<br>GGAAGTGGAATTCCGGGTGTAG<br>CGGTGAAATGCGTAGATATCCGG<br>AGGAACACCAGTGGCGAAGGC<br>GACTTCCTGGGCCAATACTGAC<br>GCTGAGGTGCGAAAGCGTGGGG<br>AGCAAACAGGATTAGATACCCT<br>GGTAGTCCACGCCGTAAACGAT<br>GAGAACTAGATGTCGGGAGAAT<br>CTGTCTTTCGGTGTCGCAGCTAA<br>CGCGTTAAGTTCTCCGCCTGGG<br>GAGTACGCCGGCAACGGTA | CM2 | 4.31 | *Woeseia* | *GU061281_s* | 97.86 | GU061281 | Uncultured bacterium clone s102 | 100.00 | AY171368.1 | marine sediment |
| TACGGGAGTGGCAAGCGTTATC<br>CGGAATTATTGGGCGTAAAGCGT<br>CCGCAGGCGGCCCTTCAAGTCT<br>GCTGTTAAAAAGTGGAGCTTAA<br>CTCCATCATGGCAGTGGAAACT<br>GTTGGGCTTGAGTGTGGTAGGG<br>GCAGAGGGAATTCCCGGTGTAG<br>CGGTGAAATGCGTAGATATCGGG<br>AAGAACACCAGTGGCGAAGGC<br>GCTCTGCTGGGCCATCACTGAC<br>GCTCATGGACGAAAGCCAGGGG<br>AGCGAAAGGGATTAGATACCCC<br>TGTAGTCCTGGCCGTAAACGATG<br>AACACTAGGTGTCGGGGGAATC<br>GACCCCCTCGGTGTCGTAGCCA<br>ACGCGTTAAGTGTTCCGCCTGG<br>GGAGTACGCACGCAAGTGTG | CM2 | 3.5 | *Synechococcus_CC9902* | *CP006882_s* | 100 | CP006882 | Uncultured bacterium clone SCG_ch08 | 100.00 | MG736619.1 | ocean subsurface |

| | | | | | | | | | | |
|---|---|---|---|---|---|---|---|---|---|---|
| TACGAAGGGACCTAGCGTAGTTCGGAATTACTGGGCTTAAAGAGTTCGTAGGTGGTTGAAAAAGTTGGTGGTGAAATCCCAGAGCTTAACTCTGGAACTGCCATCAAAACTTTTCAGCTAGAGTATGATAGAGGAAAGCAGAATTTCTAGTGTAGAGGTGAAATTCGTAGATATTAGAAAGAATACCAATTGCGAAGGCAGCTTTCTGGATCATTACTGACACTGAGGAACGAAAGCATGGGTAGCGAAGAGGATTAGATACCCTCGTAGTCCATGCCGTAAACGATGTGTGTTAGACGTTGGAAATTTATTTTCAGTGTCGCAGCGAAAGCGATAAACACACCGCCTGGGGAGTACGACCGCAAGGTTA | CM2 | 2.5 | *Clade_Ia* | *CP031125_s* | 100 | CP031125 | Candidatus Pelagibacter sp. FZCC0015 | 100.00 | CP031125.1 | seawater |
| TACGGAGGGTGCAAGCGTTGTCCGGATTTATTGGGTTTAAAGGGTGCGTAGGCGGCGTAACAAGTCAGTGGTGAAAGCCGGCAGCTCAACTGTCGAGGTGCCATTGAAACTATTATGCTTGAGTACAGACGAGGTAGGCGGAATTTATGATGTAGCGGTGAAATGCATAGATATCATAAAGAACACCGATAGCGAAGGCAGCTTACTAGGCTGTAACTGACGCTGAGGCACGAAAGCGTGGGGAGCGAACAGGATTAGATACCCTGGTAGTCCACGCTGTAAACGATGATGACTCGATGTTGGCGATAGACAGTCAGCGTCCTAGCGAAAGCGTTAAGTCATCCACCTGGGGAGTACGCTGGCAACAGTG | CM2 | 2.3 | *Cyclobacteriaceae* | *EU617868_s* | 98.92 | EU617868 | Uncultured Bacteroidetes bacterium clone T3-4 | 98.65 | KT880260.1 | microbial symbiont communities of the sun sponge |
| TACGGAGGGTGCAAGCGTTACTCGGAATCACTGGGCGTAAAGCGCATGCAGGCGGTTTAATAAGTTAGAAGTGAAATCCTACAGCTTAACTGTAGAACTGCTTCTAAAACTGTTAGACTAGAGTCTGGGAGGGGAAGATGGAATTAGTAGTGTAGGGGTAAAATCCGTAGAGATTACTAGGAATACCAAAAGCGAAGGCGATCTTCTGGAACAGTACTGACGCTGAGATGCGAAAGCGTGGGGAGCAAACAGGATTAGATACCCTGGTAGTCCACGCAGTAAACGATGAATGTTAGTCGTCGGGGCACTAGTTGTTTCGGTGATGCAGTTAACACATTAAACATTCCGCCTGGGGAGTACGGTCGCAAGATTA | CM2 | 2.2 | *Sulfurovum* | *AB278150_s* | 96.51 | AB278150 | Uncultured bacterium clone G56 | 98.39 | JX568056.1 | seafloor hydrocarbon seep |

| | | | | | | | | | | |
|---|---|---|---|---|---|---|---|---|---|---|
| TACAGAGGGTGCGAGCGTTAATCGGAATTACTGGGCGTAAAGCGCGCGTAGGCGGCTTGGTCAGTCGGATGTGAAAGCCCTGGGCTTAACCTGGGAATTGCATTCGATACTGCCAGGCTAGAATGTAGTAGAGGGAAGTGGAATTCCGGGTGTAGCGGTGAAATGCGTAGATATCCGGAGGAACATCAGTGGCGAAGGCGACTTCCTGGACTAACATTGACGCTGAGGTGCGAAAGCGTGGGGAGCAAACAGGATTAGATACCCTGGTAGTCCACGCCGTAAACGATGTCAACTAGATGTTGGGGGGTTTAACCCCTTAGTATCGCAGCTAACGCATTAAGTTGACCGCCTGGGGAGTACGGCCGCAAGGTTA | CM2 | 2.2 | *B2M28* | *HQ190975_s* | 98.93 | HQ190975 | Uncultured bacterium clone: MK0D_B22 | 99.73 | AB831353.1 | deep-sea methane-seep sediment |
| GACGAACCGTCCAAACGTTATTCGGTATCACTGGGCTTAAAGCGTGCGTAGGCGGCTTGGTAGGTGAGATGTGAAAGCCCACGGCTCAACCGTGGAATTGCGTTTCAAACCCCCAAGCTCGAGGAAGATAGGGGTGATGGGAACTTATGGTGGAGCGGTGAAATGCGTTGATATCATAGGGAACACCGGTGGCGAAAGCGCATCACTGGATCTTTTCTGACGCTGAGGCACGAAAGCTAGGGTAGCGAACGGGATTAGATACCCCGGTAGTCCTAGCCGTAAACGATGAACACTGGGTTGAGGGGACTTCCACATCCTCTCGGCCGTAGCGAAAGCGTTAAGTGTTCCGCCTGGGGAGTATGGTCGCAAGGCTG | CM2 | 2.2 | *Rubripirellula* | *Rubripirellula lacrimiformis* | 99.2 | MK559976 | Uncultured bacterium clone E133_A03 | 100.00 | KU578370.1 | ocean water |
| TACGTAGGTCCCGAACGTTGCGCGAATTTACTGGGCGTAAAGGGTCCGTAGGCGGTCTGGTAAGTGGAAGGTGAAATCCTGGGGCTCAACTCCAGAATTGCCTTCCAAACTGCTGGACTTGAGGCAGGGAGAGGTCGGCGGAATTCCCGGTGTAGCGGTGAAATGCGTAGATATCGGGAGGAACACCAGTGGCGAAGGCGGCCGACTGGAACTGTCCTGACGCTGAGGGACGAAAGCCAGGGGAGCGAACCGGATTAGATACCCGGGTAGTCCTGGCCGTAAACGATGGATGCTAGATGTGGGCAGGGAAACCTGTCCGTGTCGCAAGCTAACGCGTTAAGCATCCCGCCTGGGGAGTACGACCGCAAGGTTG | SF_G1 | 62.98% | *Hydrogenothermus* | *Hydrogenothermus marinus* | 99.2 | AJ292525 | Hydrogenothermus marinus strain VM1 | 98.93 | NR_114754.1 | Shallow water vents |

| | | | | | | | | | | |
|---|---|---|---|---|---|---|---|---|---|---|
| CACGTAGGAGGCGAGCGTTACC<br>CGGATTTACTGGGCGTAAAGCG<br>CGCGCAGGCGGCTCGGTAAGTT<br>GGGCGTGAAAGCTCCCGGCTCA<br>ACTGGGAGAGGACGTCCAAAAC<br>TGCCGGGCTAGAGGGCGGTAGA<br>GGGAGGTGGAATTCCCGGTGTA<br>GCGGTGAAATGCGTAGATATCGG<br>GAGGAACACCAGTGGCGAAGG<br>CGGCCTCCTGGACCGTCCCTGA<br>CGCTCAGGCGCGAAAGCCAGGG<br>GAGCGAACGGGATTAGATACCC<br>CGGTAGTCCTGGCCGTAAACGA<br>TGCGGACTAGGCGTTGGGCGGG<br>TCAAACCGCTCAGTGCCGTAGC<br>TAACGCGTTAAGTCCGCCGCCT<br>GGGGACTACGGCCGCAAGGCTA | SF_G1 | 8.84% | *Anaerolineaceae* | *HQ727651_s* | 91.71 | HQ727651 | Uncultured bacterium clone J10_12x-E5_0239HSNP001F_P4 | 98.66 | JN838909.1 | shallow marine hydrothermal Vent |
| TACGGAGGGTGCGAGCGTTACT<br>CGGAATTACTGGGCGTAAAGGG<br>CGCGTAGGCGGCTGGGCAAGTC<br>TGGTGTGAAAGCCCGGGGCTCA<br>ACCTCGGAAGTGCACTGGATAC<br>TGTCTGGCTTGAGTACCGGAGA<br>GGAGGGGGGAATTCCCGGTGTA<br>GCGGTGAAATGCGTAGATATCGG<br>GAGGAATACCGGTGGCGAAGGC<br>GCCCCTCTGGACGGTAACTGAC<br>GCTGAGGCGCGAAAGCGTGGG<br>GAGCAAACAGGATTAGATACCC<br>TGGTAGTCCACGCTGTAAACGAT<br>GCCCACTAGGTGTGGTGGGGGT<br>TAAGCCCTGCCGTGCCGTAGCTA<br>ACGCGTTAAGTGGGCCGCCTGG<br>GGAGTACGGCCGCAAGGTTA | SF_G1 | 4.49% | *Thermodesulforhabdus* | *Thermodesulforhabdus norvegica* | 95.43 | U25627 | Thermodesulforhabdus sp. nov. M40/2 CIV-3.2 | 99.20 | AF170420.1 | geothermally heated sediments |
| TACGGAGGTGGCGAGCGTTGCC<br>CGGAATCACTGGGCGTAAAGGG<br>GGCGTAGGCGGCCAGGCAAGTC<br>GGAGGTTAAAGCCCGGGGCTCA<br>ACCCCGGAAAGGCCTCCGATAC<br>TGCTTGGCTTGAGGGCCGGAGA<br>GGCTGGCGGAATTCCCGGTGTA<br>GGGGTGAAATCCGTAGATATCGG<br>GAGGAACACCGGTGGGGAAGC<br>CGGCCAGCTGGACGGTCCCTGA<br>CGCTGAGGCCCGAAAGCGTGGG<br>GAGCAAACCGGATTAGATACCC<br>GGGTAGTCCACGCCGTAAACGA<br>TGGGCGCTAGGTGTGGGGGGCT<br>TTATCCCTCCGTGCCGTAGCTAA<br>CGCGTTAAGCGCCCCGCCTGGG<br>GAGTACGGCCGCAAGGCTG | SF_G1 | 2.94% | *Thermosulfurimonas* | *Thermosulfurimonas dismutans* | 98.38 | LWLG01000001 | Thermosulfurimonas sp. strain F29 | 99.19 | MZ773229.1 | deep-sea hydrothermal vent |

| | | | | | | | | | | |
|---|---|---|---|---|---|---|---|---|---|---|
| TACAGAGGGTGCAAGCGTTAATCGGAATTACTGGGCGTAAAGCGCGCGTAGGTGGTTTGATAAGTTGGATGTGAAATCCCCGGGCTTAACCTGGGTCGGTCATTCAAAACTGTCAGACTAGAGTATGGTAGAGGGTAGTGGAATTTCTAGTGTAGCGGTGAAATGCGTAGATATTAGAAGGAACACCAGTGGCGAAGGCGACTGCCTGGACTGATACTGACACTGAGGTGCGAAAGCGTGGGTAGCGAACAGGATTAGATACCCTGGTAGTCCACGCCGTAAACGATGACAACTAGACGTTGGGCTCCTTAGAGGGCTTAGTGTCGAAGCTAACGCGTTAAGTTGTCCGCCTGGGGAGTACGGTCGCAAGATTA | SF_G2 | 5.2 | *endosymbionts* | *Maorithyas hadalis gill thioautotrophic symbiont I/SB3-19* | 98.93 | AB188780 | Uncultured bacterium 5133BC_bac_p1B08 | 100.00 | KT280645.1 | Hydrate Ridge methane seep anaerobic Incubation |
| TACGGGAGTGGCAAGCGTTATCCGGAATTATTGGGCGTAAAGCGTCCGCAGGCGGCCCTTCAAGTCTGCTGTTAAAAAGTGGAGCTTAACTCCATCATGGCAGTGGAAACTGTTGGGCTTGAGTGTGGTAGGGGCAGAGGGAATTCCCGGTGTAGCGGTGAAATGCGTAGATATCGGGAAGAACACCAGTGGCGAAGGCGCTCTGCTGGGCCATCACTGACGCTCATGGACGAAAGCCAGGGGAGCGAAAGGGATTAGATACCCCTGTAGTCCTGGCCGTAAACGATGAACACTAGGTGTCGGGGGAATCGACCCCCTCGGTGTCGTAGCCAACGCGTTAAGTGTTCCGCCTGGGGAGTACGCACGCAAGTGTG | SF_G2 | 3.6 | *Synechococcus_CC9902* | *CP006882_s* | 100 | CP006882 | Uncultured bacterium clone 2015-11-17-SCG_ch08 | 100.00 | MG736619.1 | Sargasso Seawater |
| TACGGAGGGTGCGAGCGTTAATCGGAATTACTGGGCGTAAAGCGCGCGTAGGCGGTTATTTAAGTCGGATGTGAAATCCCCGGGCTCAACCTGGGAACTGCATTCGATACTGGGTAACTAGAGTCTGGTAGAGGGGGGTAGAATTCCTGGTGTAGCGGTGAAATGCGTAGATATCAGGAGGAATACCAGTGGCGAAGGCGGCCCCCTGGACCAAGACTGACGCTGAGGTGCGAAAGCGTGGGGAGCAAACAGGATTAGATACCCTGGTAGTCCACGCCGTAAACGATGTCAACTAGCCGTTGGGCCCATATAAGGGTTTAGTGGCGCAGCTAACGCAATAAGTTGACCGCCTGGGGAGTACGCCGGCAACGGTA | SF_G2 | 2.95 | *Gammaproteobacteria* | *FJ517003_s* | 93.58 | FJ517003 | Uncultured bacterium clone WP12Y7D | 95.45 | KX422147.1 | "hydrothermal sulfur vent micribial Mats |

| | | | | | | | | | | |
|---|---|---|---|---|---|---|---|---|---|---|
| TACAGAGGGTGCAAGCGTTAAT<br>CGGAATTACTGGGCGTAAAGCG<br>CGCGTAGGCGGTTTGTTAAGTC<br>GGATGTGAAAGCCCCGGGCTCA<br>ACCTGGGAACTGCATTCGATACT<br>GGCAGGCTAGAGTATGGTAGAG<br>GGAAGTGGAATTCCGGGTGTAG<br>CGGTGAAATGCGTAGATATCCGG<br>AGGAACATCAGTGGCGAAGGCG<br>GCTTCCTGGACCAATACTGACGC<br>TGAGGTGCGAAAGCGTGGGGA<br>GCAAACAGGATTAGATACCCTG<br>GTAGTCCACGCCGTAAACGATG<br>AGAACTAGACGTTGGGTTCATTT<br>AAGGACTTAGTGTCGCAGCTAA<br>CGCGTGAAGTTCTCCGCCTGGG<br>GAGTACGGCCGCAAGGTTA | SF_G2 | 2.92 | *Thiogranum* | *FM179879_s* | 98.66 | FM179879 | Uncultured bacterium clone BC4 | 99.73 | JX905991.1 | shallow hydrothermal vents |
| TACGGAGGGTGCAAGCGTTAAT<br>CGGAATTACTGGGCGTAAAGCG<br>CGCGTAGGCGGCTTTGTAAGTC<br>GGATGTGAAATCCCCGGGCTCA<br>ACCCGGGAACTGCATTCGATACT<br>GCAGAACTAGAGTATGGTAGAG<br>GGAAGTGGAATTCCGGGTGTAG<br>CGGTGAAATGCGTAGATATCCGG<br>AGGAACACCAGTGGCGAAGGC<br>GACTTCCTGGGCCAATACTGAC<br>GCTGAGGTGCGAAAGCGTGGGG<br>AGCAAACAGGATTAGATACCCT<br>GGTAGTCCACGCCGTAAACGAT<br>GAGAACTAGATGTCGGGAGAAT<br>CTGTCTTTCGGTGTCGCAGCTAA<br>CGCGTTAAGTTCTCCGCCTGGG<br>GAGTACGCCGGCAACGGTA | SF_G2 | 2.35 | *Woeseia* | *GU061281_s* | 97.86 | GU061281 | Uncultured bacterium clone s102 | 100.00 | AY171368.1 | marine sediment |
| TACGAAGGGACCTAGCGTAGTT<br>CGGAATTACTGGGCTTAAAGAG<br>TTCGTAGGTGGTTGAAAAAGTT<br>GGTGGTGAAATCCCAGAGCTTA<br>ACTCTGGAACTGCCATCAAAAC<br>TTTTCAGCTAGAGTATGATAGAG<br>GAAAGCAGAATTTCTAGTGTAG<br>AGGTGAAATTCGTAGATATTAGA<br>AAGAATACCAATTGCGAAGGCA<br>GCTTTCTGGATCATTACTGACAC<br>TGAGGAACGAAAGCATGGGTAG<br>CGAAGAGGATTAGATACCCTCGT<br>AGTCCATGCCGTAAACGATGTGT<br>GTTAGACGTTGGAAATTTATTTT<br>CAGTGTCGCAGCGAAAGCGATA<br>AACACACCGCCTGGGGAGTACG<br>ACCGCAAGGTTA | SF_G2 | 2.3 | *Clade_Ia* | *CP031125_s* | 100 | CP031125 | Candidatus Pelagibacter sp. FZCC0015 | 100.00 | CP031125.1 | seawater |

| | | | | | | | | | | |
|---|---|---|---|---|---|---|---|---|---|---|
| TACGGAGGGTGCAAGCGTTAATCGGAATTACTGGGCGTAAAGCGCGCGTAGGCGGTTTGATAAGTCGGATGTGAAAGCCCTGGGCTCAACCTGGGAACTGCATTCGATACTGTCTGACTAGAGTATGGTAGAGGGAAGTGGAATTCCGGGTGTAGCGGTGAAATGCGTAGATATCCGGAGGAACATCAGTGGCGAAGGCGACTTCCTGGACCAATACTGACGCTGAGGTGCGAAAGCGTGGGGAGCAAACAGGATTAGATACCCTGGTAGTCCACGCCGTAAACGATGTCAACTAGCCGTTGGGGATATTAAAATCTTTAGTGGCGCAGCTAACGCGATAAGTTGACCGCCTGGGGAGTACGGTCGCAAGATTA | SF_G2 | 2.2 | *Thiohalophilus* | *EU652540_s* | 99.47 | EU652540 | Uncultured bacterium clone F1 NEREIS T270d | 100.00 | JF774447.1 | marine sediments microcosms |
| TACGGAGGGTCCAAGCGTTATCCGGATTTATTGGGTTTAAAGGGTCCGTAGGCGGGGTTTTAAGTCAGTGGTGAAATCCTACAGCTCAACTGTAGAACTGCCATTGAAACTGGAACTCTTGAATGTGATTGAGGTAGGCGGAATATGTCATGTAGCGGTGAAATGCTTAGATATGCCATAGAACACCGATAGCGAAGGCAGCTTACCAAGTCATAATTGACGCTGATGGACGAAAGCGTGGGGAGCGAACAGGATTAGATACCCTGGTAGTCCACGCCGTAAACGATGATCACTAGCTATTGGCGATATACAGTCAGTGGCACAGCGAAAGTGTTAAGTGATCCACCTGGGGAGTACGATCGCAAGGTTG | SF_G2 | 2.15 | *Cryomorphaceae* | *AY225660_s* | 95.95 | AY225660 | Uncultured bacterium sequence type NZ-BA-8 | 98.11 | | cirral setae of Vulcanolepas osheai |
| TACGGAGGATTCGAGCGTTATCCGGATTTATTGGGTTTAAAGGGTCCGTAGGCGGGCGATTAAGTCAGTGGTGAAATCTCACAGCTCAACTGTGAAACTGCCATTGATACTGGTTGTCTTGAATTTAGTTGAGGTGGGCGGAATACGTTATGTAGCGGTGAAATGCATAGATATAACGTAGAACACCGATTGCGAAGGCAGCTCACTAAGCTAATATTGACGCTGATGGACGAAAGCGTGGGGAGCGAACAGGATTAGATACCCTGGTAGTCCACGCCGTAAACGATGATTACTCGTTGTGCGCGATACACAGTGCGCGACTGAGCGAAAGCATTAAGTAATCCACCTGGGGAGTACGTTGGCAACAATG | GB | 7.62 | *Marinifilum* | *Marinifilum fragile* | 100 | BAZX01000063 | Marinifilum fragile CECT 7942 strain JC2469 | 100.00 | NR_044597.2 | tidal flat Sediment |

| | | | | | | | | | | |
|---|---|---|---|---|---|---|---|---|---|---|
| TACGTAGGGAGCAAGCGTTGTCCGGATTTACTGGGTGTAAAGGGCGCGCAGGCGGGTTGGTAAGTCAGAGGTGAAATCCTACAGCTTAACTGTAGAACTGCCTTTGATACTGCTGATCTTGAGTATGGAAGAGAGAGACGGAATTCCAGGTGTAGTGGTGAAATACGTAGATATCTGGAAGAACACCAGTTGCGAAGGCGGTCTCTTGGTCCAATACTGACGCTGAGGCGCGAAAGCGTGGGTAGCAAACAGGATTAGATACCCTGGTAGTCCACGCTGTAAACGATGAATACTAGGTGCTGGGTCTTTAGATTCAGTGTCGCAGCTAACGCATTAAGTATTCCACCTGGGGAGTACGATCGCAAGGTTG | GB | 5.87 | *PHOS-HE36* | *JQ580230_s* | 98.92 | JQ580230 | Uncultured bacterium clone APC-3439-J3F12 | 99.46 | KF616740.1 | Hydrate Ridge |
| TACGGAGGGGGCAAGCGTTATCCGGAATCACTGGGCGTAAAGAGCGCGTAGGCGGGTTAAAAAGTCGGGCGTGAAATTTATCGGCTTAACTGATAAATGTCGTCCGATACTTTTAATCTTGAGGATAGGAGAGGAGAGTAGAATTCCCGGTGTAGCGGTGAAATGCATTGATATCGGGAGGAATGCCAGTTGCGAAGGCGGCTCTCTGGAATATTCCTGACGCTGAGGCGCGAAAGCGTGGGTATCGAACCGGATTAGATACCCGGGTAGTCCACGCCGTAAACGATGGATGTTAGGTGTAGGGGGTTACTCCTGTGCCGTAGCTAACGCGTTAAACATCCCGCCTGGGGAGTACGGTCGCAAGGCTG | GB | 3.28 | *NA* | *ASOY_s* | 83.15 | ASOY01000050 | Uncultured bacterium clone 490CT10B38 | 84.55 | KX953010.1 | deep sea sediment |
| TACGGAGGGTGCAAGCGTTGTCCGGATTTATTGGGTTTAAAGGGTGCGTAGGCGGCGTAACAAGTCAGTGGTGAAAGCCGGCAGCTCAACTGTCGAGGTGCCATTGAAACTATTATGCTTGAGTACAGACGAGGTAGGCGGAATTTATGATGTAGCGGTGAAATGCATAGATATCATAAAGAACACCGATAGCGAAGGCAGCTTACTAGGCTGTAACTGACGCTGAGGCACGAAAGCGTGGGGAGCGAACAGGATTAGATACCCTGGTAGTCCACGCTGTAAACGATGATGACTCGATGTTGGCGATAGACAGTCAGCGTCCTAGCGAAAGCGTTAAGTCATCCACCTGGGGAGTACGCTGGCAACAGTG | GB | 2.53 | *Cyclobacteriaceae* | *EU617868_s* | 98.92 | EU617868 | Uncultured Bacteroidetes bacterium clone T3-4 | 98.65 | KT880260.1 | tidal creek sediment |

| | | | | | | | | | | |
|---|---|---|---|---|---|---|---|---|---|---|
| CACGGGGGGGGCGAGCGTTATTCGGAATTACTGGGCGTAAAGGGCGCGTAGGCGGTCCGTTAAGTGTGAAGTGAAATGCCTGGGCTCAACCTGGGACGTGCTTTGCATACTGGTGGACTTGAGTCCAAGAGGGGGTGGTGGAATTCCTGGTGTAGGGGTGAAATCCGTAGATATCAGGAGGAACACCGTTGGCGAAGGCGGCCACCTGGATTGGTACTGACGCTGAGGCGCGAAAGCGTGGGGAGCGAACAGGATTAGATACCCTGGTAGTCCACGCCGTAAACGATGTTCACTTGGTGTTGGTGTGATTAACCACATCAATGCCGGAGCTAACGCATTAAGTGAACCGCCTGGGGAGTACGGTCGCAAGGCTG | GB | 2.53 | *Thermotomaculum* | *HQ916584_s* | 95.19 | HQ916584 | Uncultured bacterium clone FS396_454 | 100.00 | DQ909392.1 | marine hydrothermal vent fluids |
| TACGTAGGGGGCAAGCGTTGTCCGGATTTACTGGGTGTAAAGGGCGCGTAGGCGGGTTTGTAAGTCAGTGGTGAAATCCTACAGCTTAACTGTAGAACTGCCTTTGATACTGCAGACCTTGAGTACGGAAGAGAGAGGCGGAATTCCAGGTGTAGTGGTGAAATACGTAGATATCTGGAAGAACACCAGAGGCGAAGGCGGTCTCTTGGTCCGTTACTGACGCTGAGGCGCGAAAGCGTGGGGAGCAAACAGGATTAGATACCCTGGTAGTCCACGCCGTAAACGATGAATACTAGGTGTTGGGTTTTTAACTCAGTGCCGCAGCAAACGCATTAAGTATTCCACCTGGGGAGTACGATCGCAAGGTTG | GB | 2.17 | *IheB3-7* | *JQ579954_s* | 97.83 | JQ579954 | Uncultured bacterium clone D1-STK-23 | 98.37 | MH091110.1 | mangrove sediments |
| TACGGGGGGTGCAAGCGTTATTCGGATTTACTGGGCGTAAAGCGCGCGTAGGCGGCCGTTTAAGTCAGATGTGAAAGCCCGGGGCTCAACCCCGGAAGTGCATTTGATACTATTCGGCTTGAGTATGGGAGAGGGAAGTGGAATTCCTGGTGTAGAGGTGAAATTCGTAGATATCAGGAGGAACACCGGTGGCGAAGGCGACTTCCTGGACCAATACTGACGCTGAGGCGCGAAGGCGTGGGGAGCAAACAGGATTAGATACCCTGGTAGTCCACGCAGTAAACGGTGATCACTAGGTGTAGCGGGTATTGACCCCTGCTGTGCCGCAGCTAACGCATTAAGTGATCCGCCTGGGGAGTACGGCCGCAAGGTTA | GB | 2.1 | *Desulfosarcinaceae* | *FM242222_s* | 99.73 | FM242222 | Uncultured bacterium clone C158 | 100.00 | KJ817702.1 | soil |

| | | | | | | | | | | |
|---|---|---|---|---|---|---|---|---|---|---|
| TACCCGCGGTCCAAGTCGCAGCCATTTTTATTGGGTCTAAAACATCCGTAGCTTGCTCTTTAAGTTCCTTGTGAAATCCTATATCTTAAATATAGGGCGTGCAGGGAATACTACTGAGCTAGAGACTGGAAGACGTAACGAGTACGTTCGAAGTAGCGGTTAAATGTGTTAATCTCGAGCGGACTAACAATAGCGAAGGCACGTTACGAGGACAGTTCTGACAGTAAGGGATGAAGGCTAGGGGCGCAAAACGGATTAGATACCCGTGTAGTCCTAGCAGTAAACACTGTACACTAAACATTAGTACCTCCTCGAGAGGTATTGGTGCTGAAGCGAAGGCGAAGAGTGTACTACCTGGGAAGTATAGTCGCAAGGCCG | GB | 2.07 | *SCGC_AAA011-D5* | *EU731474_s* | 94.18 | EU731474 | Uncultured archaeon O127906H12 | 94.97 | FN865760.1 | NA |
| TACGGAGGGTGCAAGCGTTATCCGGATTCATTGGGTTTAAAGGGTGCGCAGGCGGACTTTTAAGTCAGTGGTGAAATCCCGGGGCTCAACCCCGGAACTGCCATTGATACTGAAAGTCTTGAGTTTGGTTGAAGTAGGCGGAATGTAGCATGTAGCGGTGAAATGCTTAGATATGCTACAGAACACCGATCGCGAAGGCAGCTTACTAAACCAATACTGACGCTCAGGCACGAAAGCGTGGGGAGCGAACAGGATTAGATACCCTGGTAGTCCACGCCGTAAACTATGATCACTCGTTGTTGGCGATACACAGTCAGCGACCAAGCGAAAGCGATAAGTGATCCACCTGGGGAGTACGATCGCAAGGTTG | SF_EF | 5 | *Bacteroidetes_VC2.1_Bac22* | *AY197392_s* | 98.87 | AY197392 | Uncultured Cytophaga sp BJS72-056 | 100.00 | AB238986.1 | cold seep sediment |
| TACGGAGGATGCAAGCGTTATCCGGATTTATTGGGTTTAAAGGGTGCGCAGGCGGCCTTATAAGTCAGTGGTGAAATCTCTCGGCTCAACCGAGAAACTGCCATTGATACTGTAGGGCTAGAATACAGACGAGGTAGGCGGAATGTAGCATGTAGCGGTGAAATGCTTAGATATGCTACAGAACACCGATCGCGAAGGCAGCTTACCAGGCTGTTATTGACGCTAATGCACGAAAGCGTGGGGAGCGAACAGGATTAGATACCCTGGTAGTCCACGCCGTAAACGATGATCACTCGATGTTGGCGATAGACAGTCAGCGTCCAAGCGAAAGTATTAAGTGATCCACCTGGGGAGTACGATCGCAAGGTTG | SF_EF | 3.81 | *Lentimicrobiaceae* | *FJ497390_s* | 99.46 | FJ497390 | Uncultured Bacteroidetes bacterium clone VS_CL-139 | 99.46 | FJ497390.1 | Vailulu'u Seamount |

| | | | | | | | | | | |
|---|---|---|---|---|---|---|---|---|---|---|
| TACGGGGGGAGCAAGCGTTGTCCGGAATTACTGGGCGTAAAGGGCGTGTAGGTGGGCTGATAAGTCAGATGTGAAAGCCCGCGGCTTAACCGCGGAACTGCATTTGAAACTGTCAGTCTTGAGTACGAGAGAGGGTAGTGGAATTCCCAGTGTAGCGGTGAAATGCGTAGATATTGGGAAGAACACCAGTAGCGAAGGCGGCTACCTGGCTCGCAACTGACGCTAATGCGCGAAAGCGTGGGGAGCAAACAGGATTAGATACCCTGGTAGTCCACGCTGTAAACGATGGGCACTAGGTGTCGGTTCCGCTTGCGGAATCGGTGCCGCAGCAAACGCATTAAGTGCCCCGCCTGGGGAGTACGATCGCAAGGTTG | SF_EF | 3.33 | *Caldithrix* | *AY280423_s* | 94.1 | AY280423 | Uncultured bacterium Acs2P47 | 98.66 | AB292968.1 | hydrothermal sulfide structure |
| TACGTAGGGGGCAAGCGTTGTCCGGATTTACTGGGTGTAAAGGGCGCGTAGGCGGGTTTGTAAGTCAGTGGTGAAATCCTACAGCTTAACTGTAGAACTGCCTTTGATACTGCAGACCTTGAGTACGGAAGAGAGAGGCGGAATTCCAGGTGTAGTGGTGAAATACGTAGATATCTGGAAGAACACCAGAGGCGAAGGCGGTCTCTTGGTCCGTTACTGACGCTGAGGCGCGAAAGCGTGGGGAGCAAACAGGATTAGATACCCTGGTAGTCCACGCCGTAAACGATGAATACTAGGTGTTGGGTTTTTAACTCAGTGCCGCAGCAAACGCATTAAGTATTCCACCTGGGGAGTACGATCGCAAGGTTG | SF_EF | 3.11 | *IheB3-7* | *JQ579954_s* | 97.83 | JQ579954 | Uncultured bacterium clone D1-STK-23 | 98.37 | MH091110.1 | mangrove sediments |
| TACGGAGGGTGCAAGCGTTAATCGGAATCACTGGGCGTAAAGCGTGCGTAGGCTGCGCTTCAAGTCAGACGTGAAAGCCCTCGGCTCAACCGAGGAATTGCGTTTGAAACTGGAGTGCTTGAGTCTCGGAGAGGTTGGCGGAATTCCTGGTGTAGGAGTGAAATCCGTAGATATCAGGAGGAACACCGGCGGCGAAGGCGGCCAACTGGACGAGTACTGACGCTGAGGTACGAAAGCGTGGGTAGCAAACAGGATTAGATACCCTGGTAGTCCACGCTGTAAACGATGGATATTAGGTGTCGGGGTTCACACTTCGGTGCCGCAGTTAACGCGTTAAATATCCCGCCTGGGGAGTACGGTCGCAAGGCTG | SF_EF | 2.72 | *Desulfovibrio* | *Pseudodesulfovibrio nedwellii* | 97.56 | LC752232 | Pseudodesulfovibrio sp. SYK | 97.30 | LC752232.1 | saline lake sediment |

| | | | | | | | | | | |
|---|---|---|---|---|---|---|---|---|---|---|
| TACCGGCGGCTCGAGTGGTGGCCGCTATTACTGGGCTTAAAGCGTCCGTAGCTTGGTCGTTAAGTCTCTGGGGAAATCTTCCGGCTCAACCGGAAGGCGTCTCAGGGATACTGGCGGCCTAGGGATCGGGAGAGGTGAGAGGTACTCTGGGGGTAGGAGTGAAATCCTGTAATCCTCAGGGGACCACCTGTGGCGAAGGCGTCTCACCAGAACGACTCCGACAGTGAGGGACGAAAGCTGGGGGAGCAAACCGGATTAGATACCCGGGTAGTCCCAGCCGTAAACGATGCGCGTTAGGTGTATCGGTGACCACGAGTTACCGAGGTGCCGAAGGGAAACCGTGAAACGCGCCGCCTGGGAAGTACGGTCGCAAGGCTG | SF_EF | 2.71 | *Methanofollis* | *Methanofollis fontis* | 99.47 | MG437305 | Methanofollis fontis strain FWC-SCC2 | 99.47 | MG437305.1 | Methane marine sediment |
| TACGGAGGGTGCAAGCGTTACTCGGAATCACTGGGCGTAAAGCGCATGCAGGCGGTTTAATAAGTTAGAAGTGAAATCCTACAGCTTAACTGTAGAACTGCTTCTAAAACTGTTAGACTAGAGTCTGGGAGGGGAAGATGGAATTAGTAGTGTAGGGGTAAAATCCGTAGAGATTACTAGGAATACCAAAAGCGAAGGCGATCTTCTGGAACAGTACTGACGCTGAGATGCGAAAGCGTGGGGAGCAAACAGGATTAGATACCCTGGTAGTCCACGCAGTAAACGATGAATGTTAGTCGTCGGGGCACTAGTTGTTTCGGTGATGCAGTTAACACATTAAACATTCCGCCTGGGGAGTACGGTCGCAAGATTA | SF_EF | 2.5 | *Sulfurovum* | *AB278150_s* | 96.51 | AB278150 | Uncultured bacterium clone G56 | 98.39 | JX568056.1 | seafloor hydrocarbon seep |
| TACGGAGGATGCAAGCGTTATCCGGATTTATTGGGTTTAAAGGGTACGTAGGCGGAAAATTAAGTCAGTAGTGAAATCCTGCAGCTTAACTGTAGAACTGTTATTGATACTGGTTTTCTTGAATATAGTTGAGGTAGGCGGAATGTGTAATGTAGCGGTGAAATGCTTAGATATTACACAGAACACCGATTGCGAAGGCAGCTTACTAAGCTATGATTGACGCTGAGGTACGAAAGCGTGGGGAGCGAACAGGATTAGATACCCTGGTAGTCCACGCCGTAAACGATGATCACTCGTTGTTGGCAATACATCGTCAGCGACTGAGCGAAAGCATTAAGTGATCCACCTGGGGAGTACGCTCG | SF_EF | 2.1 | *Bacteroidetes_BD2-2* | *JF344511_s* | 99.19 | JF344511 | Uncultured Bacteroidales bacterium clone SD08_034 | 100.00 | KJ566252.1 | sediment |

| |
|---|
| CAAGAGTG |

**Supplementary table 2**. Envfit results against nMDS1, nMDS2- Jaccard weighted. One star (*); p-value less than 0.01, two stars (**) p-value is less than 0.001. Only variables with a p-value of less than 0.05 are marked in bold and considered statistically significant.

| Variable | nMDS1 | nMDS2 | $r^2$ | Pr (>r) | |
|---|---|---|---|---|---|
| temp | 0.92386 | 0.38272 | 0.4129 | 0.1113 | |
| **psu** | 0.40980 | 0.91218 | 0.8374 | 0.0018 | ** |
| do | 0.90785 | 0.41929 | 0.1342 | 0.5714 | |
| orp | 0.54389 | 0.83916 | 0.3899 | 0.1254 | |
| ph | -0.05997 | -0.99820 | 0.2043 | 0.3986 | |
| **ms.cm** | 0.44242 | 0.89681 | 0.7311 | 0.0135 | * |
| mVph | 0.20780 | 0.97817 | 0.3928 | 0.1389 | |
| **ppm.tds** | 0.44342 | 0.89631 | 0.7342 | 0.0134 | * |
| **Cl** | 0.42823 | 0.90367 | 0.7001 | 0.0159 | * |
| **Br** | 0.42841 | 0.90358 | 0.6867 | 0.0173 | * |
| SO4 | 0.62616 | 0.77969 | 0.3819 | 0.1260 | |
| **Na** | 0.41695 | 0.90893 | 0.7149 | 0.0159 | * |
| NH3 | 0.30526 | 0.95227 | 0.3409 | 0.1890 | |
| **K** | 0.65368 | 0.75677 | 0.7953 | 0.0314 | ** |
| **Mg** | 0.31056 | 0.95055 | 0.7771 | 0.1161 | ** |
| Ca | 0.91023 | 0.41411 | 0.1880 | 0.2997 | |
| Mn | -0.54906 | 0.83578 | 0.3061 | 0.2246 | |
| Fe | -0.60695 | 0.79474 | 0.3251 | 0.1971 | |
| Zn | -0.36158 | -0.93234 | 0.1583 | 0.4973 | |
| As | 0.76016 | 0.64974 | 0.3356 | 0.1860 | |
| Rb | -0.06609 | 0.99781 | 0.2447 | 0.3202 | |
| **Sr** | 0.62092 | 0.78388 | 0.6330 | 0.0201 | * |
| Mo | 0.74979 | 0.66168 | 0.1684 | 0.4646 | |
| Cs | 0.90667 | 0.42183 | 0.3881 | 0.1342 | |

**Supplementary table 3**. Envfit results against nMDS1, nMDS2- Jaccard unweighted. One star (*); p-value less than 0.05, two stars (**) p-value is less than 0.01. Only variables with a p-value of less than 0.05 are marked in bold and considered statistically significant.

| Variable | nMDS1 | nMDS2 | $r^2$ | Pr (>r) | |
|---|---|---|---|---|---|
| temp | 0.84990 | 0.52695 | 0.4100 | 0.1257 | |
| **psu** | 0.99650 | 0.08361 | 0.5575 | 0.0380 | * |
| do | 0.28411 | 0.95879 | 0.2883 | 0.2515 | |
| orp | 0.69207 | 0.72183 | 0.3233 | 0.2077 | |
| ph | -0.56141 | 0.82754 | 0.4068 | 0.1282 | |
| ms.cm | 0.98627 | 0.16512 | 0.5031 | 0.0613 | |
| mVph | 0.73503 | -0.67803 | 0.4937 | 0.0697 | |
| ppm.tds | 0.98467 | 0.17444 | 0.5044 | 0.0612 | |
| Cl | 0.99286 | 0.11930 | 0.4792 | 0.0739 | |
| Br | 0.99393 | 0.10997 | 0.4719 | 0.0796 | |
| SO4 | 0.92180 | 0.38765 | 0.3248 | 0.2007 | |
| Na | 0.99463 | 0.10350 | 0.4827 | 0.0752 | |
| K | 0.79606 | 0.60522 | 0.7080 | 0.1721 | |
| Mg | 0.90212 | 0.43149 | 0.3885 | 0.1850 | |
| Ca | 0.13836 | 0.99038 | 0.6906 | 0.9484 | |
| Mn | 0.18578 | -0.98259 | 0.4590 | 0.0824 | |
| Fe | 0.14126 | -0.98997 | 0.4838 | 0.0653 | |
| Zn | -0.56923 | -0.82218 | 0.1064 | 0.6503 | |
| As | 0.86097 | 0.50866 | 0.3213 | 0.2107 | |
| Rb | 0.53325 | -0.84596 | 0.3308 | 0.2023 | |
| **Sr** | 0.99571 | 0.09249 | 0.5824 | 0.0303 | * |
| Mo | 0.42101 | 0.90706 | 0.2559 | 0.3066 | |
| Cs | 0.88616 | 0.46339 | 0.3857 | 0.1425 | |
